# The N-terminal extension region of BamE coordinates interaction between the β-barrel domain of BamA and BamD for efficient assembly

**DOI:** 10.1101/2024.10.31.621230

**Authors:** Yuki Maruno, Edward Marshall Germany, Yukari Nakajima, Rei Ohyama, Yukihiko Masumura, Takuya Shiota

## Abstract

In the Gram-negative bacteria, the assembly of β-barrel proteins into the outer membrane is mediated by the β-barrel Assembly Machinery (BAM) complex. BamA is a central component of the BAM complex. The molecular mechanisms of BamA when the substrate protein is assembled has been well elucidated and further the inhibitors targeting key mechanisms of BamA have been isolated. Although the BAM complex is a multi-subunit machine and its coordination is necessary for full function, the molecular mechanism remains unclear. Here we report on a fine-tuning system with a subunit called BamE, which is required for the cooperation of two essential subunits BamA and BamD. BamE is a lipoprotein with an N-terminal extension region between the lipid-modifying cysteine on the N-terminus and the periplasmic folding domain. Our interaction mapping revealed that the N-terminal extension region of BamE interacts with the β-barrel domain of BamA, which contains a catalytic site for assembly termed lateral gate. The mutational analysis demonstrated that the N-terminal extension region of BamE fine-tunes the assembly stage dependent BamA-BamD interaction, responsible for the crucial role for assembly and integrity of the outer membrane. We suggest that this fine-tuning system plays an important role in the substrate relay system from the receptor BamD to the insertase BamA. Furthermore, structure prediction-based analysis suspects that the C-terminal region of BamD acts as the replacement of BamE in BamE-less bacteria.

## Introduction

The outer membrane (OM), a key feature of Gram-negative bacteria, consists of phospholipids, lipopolysaccharide, outer membrane proteins (OMPs), and lipoproteins (Bos and Tommassen, 2004). OMPs are the principal component of the OM and responsible for various essential biological reactions such as transport of nutrients, toxin secretion, and acting as efflux pumps (Guest and Silhavy, 2023; Hayashi et al., 2024a). OMPs are characterized by the presence of a β-barrel transmembrane domain anchoring the proteins in the OM (Hayashi et al., 2024a). The essential biosynthesis process of OMPs is assembly: insertion of the β-barrel transmembrane domain into the OM with proper folding (Guest and Silhavy, 2023; Hayashi et al., 2024b).

The assembly of most OMPs is performed by the β-barrel Assembly Machinery (BAM) complex (Guest and Silhavy, 2023; Tomasek and Kahne, 2021). In *Escherichia coli*, the BAM complex is a heteropentamer formed by BamA-E subunits. BamA, the core subunit, is an OMP itself, while BamB-E are lipoproteins (Heinz and Lithgow, 2014; Sklar et al., 2007; Tomasek and Kahne, 2021).The BamA and BamD subunits are essential for *E. coli* viability, with BamB, C, E functioning as accessory proteins. BamA is conserved in all Gram-negative bacteria as Omp85 superfamily proteins, with homologues identified in eukaryotic cells as Sam50, a mitochondria; OEP80, in chloroplasts (Hayashi et al., 2024b; Heinz and Lithgow, 2014). The BamD subunit is widely conserved in Gram-negative bacterial lineages (Anwari et al., 2012). While BamB and BamE are conserved in alpha, beta, and gamma-proteobacteria, BamC is conserved in beta and gamma, but not alpha. In delta and epsilon-proteobacteria, homologues of these accessory proteins have not been identified (Anwari et al., 2012).

BamA plays a central role in assembling the newly synthesized OMPs (substrates) (Doyle and Bernstein, 2022; Tomasek and Kahne, 2021). During the assembly of newly synthesized OMPs, the first and last β-strands of the β-barrel transmembrane region, forming the “lateral gate”, of BamA change conformation dynamically (Noinaj et al., 2014; Tomasek et al., 2020). The lateral gate recognizes a β-signal in the most C-terminal β-sheet and catalyzes the formation of the substrate OMP β-sheet (Doyle et al., 2022; Tomasek et al., 2020). The N-terminal region of BamA is exposed into the periplasmic region and folds five repeats of the polypeptide transport-related (POTRA) domains (Noinaj et al., 2013; Voulhoux et al., 2003). The POTRA domains are numbered from the N-terminus as POTRA 1 to POTRA 5. POTRA 2 and POTRA 5 bind both N-terminal and C-terminal sides of BamD, respectively, and thus form a funnel-like structure (Bakelar et al., 2016; Gu et al., 2016; Iadanza et al., 2016). Accessory subunits dock outside of the funnel-like structure. BamD facilitates the partial folding (pre-folding) of the substrate inside the funnel-like structure before insertion at the lateral gate (Doyle et al., 2022). Recent study revealed that the substrates contain the signal sequence inside of the protein termed “internal signal” as well as the β-signal and the dual recognition of β-signal and internal signals by BamD is important for assembly (Germany et al., 2024).

The assembly of substrate requires the cooperation of pre-folding recognition by BamD and insertion by the lateral gate of BamA (Hagan et al., 2015; Hart et al., 2020). The lateral gate of BamA directly connects to POTRA 5, which interacts with BamD (McCabe et al., 2017a). BamE is located in the vicinity of POTRA 5-BamD binding site and stabilizes the interaction of POTRA 5 and BamD (Bakelar et al., 2016; Gu et al., 2016; Iadanza et al., 2016). A recent study demonstrated that BamE has independent specific binding regions for POTRA 5 and BamD, respectively (Kumar and Konovalova, 2023). The interactions through these regions facilitated the stability of BamA-BamD binding and the efficiency of the substrate OMP assembly. Another study demonstrated that simply restoring the BamA-BamD binding lost by mutation by further mutation does not restore the assembly function, implying that BamA POTRA 5 and BamD should be dynamically changed, not simply stably bound (McCabe et al., 2017a; Ricci et al., 2012). Multiple BAM complex structures reported in this decade have shown that the periplasmic domain of the BAM complex takes several states, also supporting its dynamics (Bakelar et al., 2016; Fenn et al., 2024; Gu et al., 2016; Iadanza et al., 2016; Noinaj et al., 2013). Our recent *in vivo* study demonstrated that the BamE deletion does not completely dissociate BamA and BamD, but affects partial binding, which is required for assembly (Thewasano et al., 2023). Moreover, BamE modulates the conformation of BamA, likely through its interactions with BamD (Rigel et al., 2012). These findings imply that BamE, along with stabilizing BamA-D binding, fine-tunes the dynamic nature of this interaction; however, the molecular mechanism of how BamE achieves the fine-tuning of dynamics of BamA-D interaction remains unclear.

In this study, we built an *in vivo* interaction map of BamE using *in situ* photo-crosslinking to understand the fine-tuning system of BamA-BamD binding by BamE. *In situ photo-crosslinking* employs the photo-reactable unnatural amino acid, which is the *p*-benzoyl-L-phenylalanine (*p*BPA) to capture protein-protein proximity by forming a covalent bond of *p*BPA with nearby protein *in vivo* (Chin et al., 2002). This method can exclude artificial effects such as detergent solubilization, as the interaction of proteins *in vivo* can be assessed more directly. Interestingly, we show the N-terminal region of BamE interacts in the vicinity of the periplasmic boundary of the BamA β-barrel domain. This region contains a highly conserved tyrosine residue, Y28. Our mutational analysis demonstrated that Y28 assists in the proximity of BamD to the transmembrane domain of BamA, important for substrate assembly but not the stability of the BamA-BamD binding. Our findings revealed that BamE stabilizes the binding of BamD to POTRA 5 of BamA in the C-terminal region and regulates the distance between the transmembrane domains of BamD and BamA in the N-terminal region, thereby fine-tuning the linkage between BamA and BamD during assembly.

## Results

### BamE is responsible for the stability and proper positioning of BamA-BamD binding

So far, the impact of the loss of BamE to BamA-BamD binding has been evaluated through the stability of the detergent solubilized complex (Kumar and Konovalova, 2023; McCabe et al., 2017b; Ricci et al., 2012). Previous reports showed that the loss of BamE resulted in the complete dissociation of BamD from BamA in detergent-solubilized BAM complex. However, cell growth of the bamE deletion showed more moderate than expected from the dissociation of two essential components of the BAM complex, in other words, the loss of one of the essential components of the BAM complex. Our recent report using *in situ* photo cross-linking showed that even bamE deletion *E. coli* retained the proximity of the BamD to BamA, indicating that the evaluation of BamE impact by detergent solubilization overestimates than the actual effect *in vivo* (Thewasano et al., 2023).

To analyze the exact contribution of BamE in BamA-BamD binding, we built a map of the sites where BamA interacts with BamD or BamE *in vivo* by *in situ* photo cross-linking. We introduced *p*BPA at 95 different residues of BamA. To ensure that *p*BPA-introduced BamA to preferentially form the BAM complex with other subunits, we used a BamA shut-off strain which can regulate the expression of endogenous BamA (Lehr et al., 2010). Also, BamA and its cross-linked products were purified by introducing the hexahistidine tag into the downstream of the SEC signal sequence of BamA, and sufficient cross-linked products were prepared by concentrating the eluted fractions. UV-dependent cross-linked products were assessed by immunoblotting using anti-BamA, anti-BamD, and anti-BamE antibodies (Fig. S1, S2, and S3) (Gunasinghe et al., 2018). BamA-BamD cross-linked products were detected at 15 different positions in POTRA domains 1, 2, and 5, which is consistent with previous studies (Fig. 1 A) (Gu et al., 2016; Thewasano et al., 2023). Interestingly, BamA-BamD interactions were observed in POTRA 3 and POTRA 4, and within the β-barrel domain (turn 1, turn 2, turn 5, and β-strand 16). These positions did not correspond to structures of the BAM complex. BamE also interacted not only with POTRA5 but also with turn 2 of BamA.

**Figure 1.**
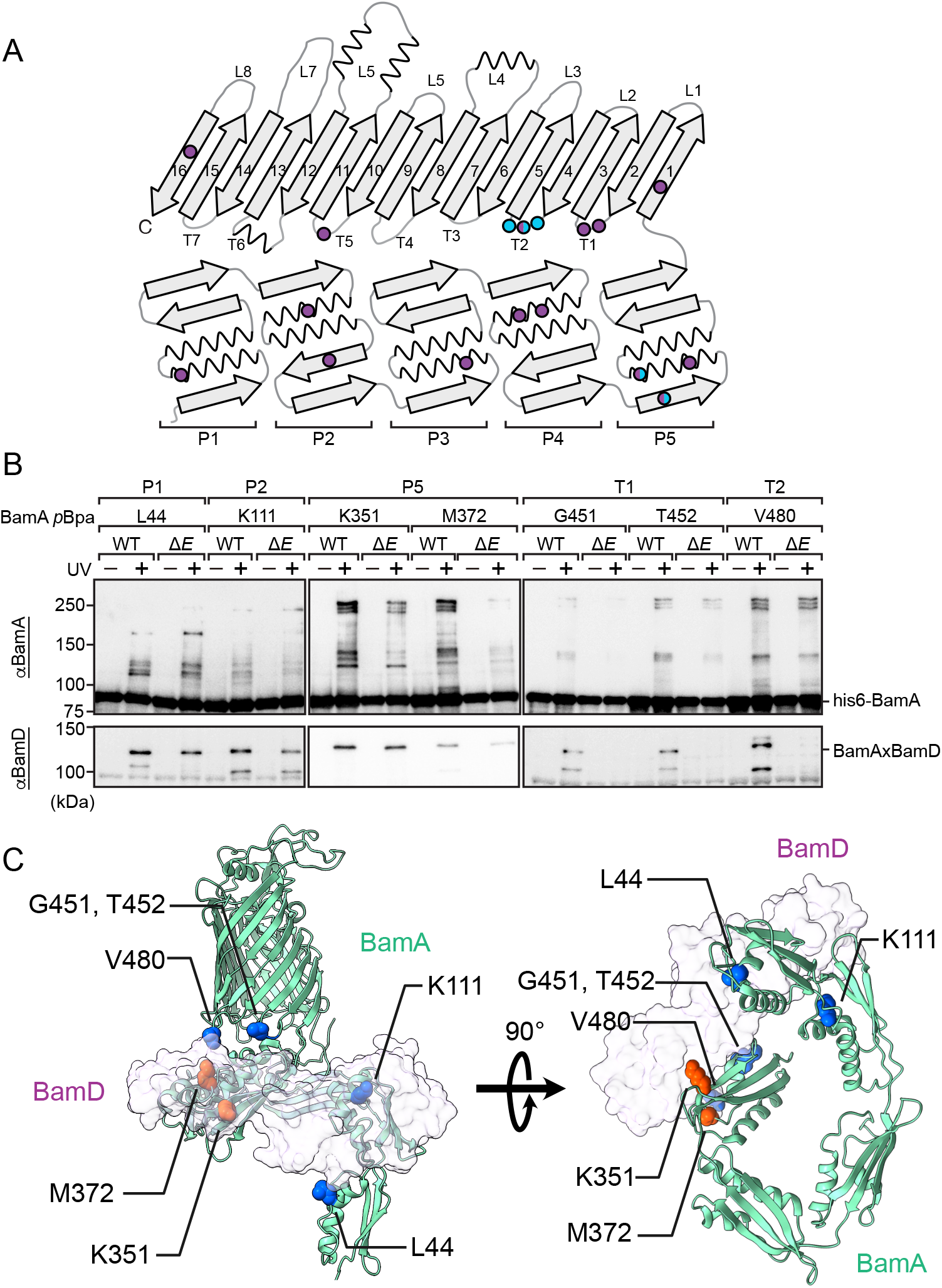
BamE is required for proper BamA-BamD interactions. (A) Interaction map of BamA on its schematic secondary structure. Cross-linked positions with BamD and BamE were indicated in purple and sky blue, respectively. (B) *In situ* photo-crosslinking analysis of *p*BPA at the indicated positions of BamA. BamA-BamD cross-linked products were compared with BL21 (WT) and Δ*bamE* strains. Cross-linked products were detected with anti-BamA antisera (top) or anti-BamD antisera (bottom). Immunoblots are representative of at least two biological replicates. (C) Mapping of BamA-BamD cross-linked residues onto the structure of BAM complex (5AYW). BamA and BamD are shown by ribbon and transparent surface model, respectively. Bottom view, excluding the β-barrel domain, is shown on the right and side view is shown on the left. The positions of BamA where the BamA-BamD cross-linked products were reduced in Δ*bamE* strain compared to WT strain are indicated in blue, the positions of BamA that did not change are indicated in red.

We then repeated *in situ* photo cross-linking at seven representative positions of BamA (residues 44, 111, 351, 372, 451, 452, and 480) in the BL21 or Δ*bamE* strains to analyze the effect of BamE on site-specific BamA-BamD binding (Fig. 1 B). While BamA-BamD cross-linked products at periplasmic regions of the BAM complex, POTRA 1 (44), POTRA 2 (111), POTRA 5 (351 and 372) were slightly decreased in Δ*bamE*, the cross-linked products at β-barrel region, turn 1 (451 and 452), and turn 2 (480), disappeared depending on the lack of the BamE (Fig. 1 B and C). Previous neutron reflectometry analysis demonstrated that BamE changes the conformation of the POTRA domain of BamA (Chen et al., 2021; Rigel et al., 2012; Thewasano et al., 2023). Taken together with our findings BamE stabilizes the BamA-BamD binding and regulates the correct positions of BamD relative to BamA.

### The N-terminal extension region of BamE is important for substrate assembly

To reveal the molecular mechanisms of regulation of BamA-BamD binding by BamE, we further built an interaction map of BamE *in vivo*. We expressed C-terminal hexahistidine tagged BamE harboring *p*BPA in the Δ*bamE* strain and the cross-link products were purified with Ni-NTA as same as BamA cross-linking (Thewasano et al., 2023). We confirmed that attaching the hexahistidine tag to the C-terminal of BamE did not impair the BamE function by directly analysing OMP assembly *in vitro* reconstitution methods (Fig. S4 A and B). The *p*BPA was introduced at 20 different positions of BamE, respectively (Fig. 2 A and B). Crosslink products with BamA or BamD were formed at 13 positions (P30, N41, V48, Y57, L63, M64, F68, G69, R78, H83, V86, N103, and N106) and seven positions (L63, M64, F68, R78, T87, T92, and K107), respectively. These crosslink products were generally consistent with the structures and important residues for functions in previous reports. For example, N41, or L63, M64, and F68 were in close structurally proximity to Y37, an amino acid previously reported to be important for interaction with BamA, or D66, an important residue for interaction with BamD, respectively (Kumar and Konovalova, 2023).

Since BamE is a lipoprotein, an unfolded N-terminal extension region connects the C20, the lipid-modifying residue, to the periplasmic domain (I32 to K107). The P30, the position prominently cross-linked to BamA, was in the N-terminal extension region of BamE (Fig. 2 A and B). Previous reported structure showed that the N-terminal extension region was close to the turn 2 in the β-barrel domain of BamA (Bakelar et al., 2016; Doyle et al., 2022; Fenn et al., 2024; Gu et al., 2016; Han et al., 2016; Haysom et al., 2023; Iadanza et al., 2016; Kaur et al., 2021; Miller et al., 2022; Seyfert et al., 2023; Shen et al., 2023; White et al., 2021; Wu et al., 2021; Xiao et al., 2021). In our mapping of BamA in the present study, the V480*p*BPA of turn2 was cross-linked to BamE, which is also consistent with this finding (Fig. 1 A). Analysis of sequence conservation of the N-terminal extension region of BamE revealed that tyrosine 28 was highly conserved (Fig. 2 C). In a previous report, the importance of a specific residue in BamE was identified by mutation of the particular residues with glycine. The significance of this residue was determined based on the vancomycin resistance of point mutants, which reflects the integrity of the OM (Knowles et al., 2011). Following this approach, we constructed the point mutant Y28G to examine whether the proximity of the N-terminal extension region of BamE and turn 2 of BamA is important or not. We also tested M64G and D66G where the binding region BamD previously reported (Fig. 3 A) (Kumar and Konovalova, 2023). We transformed the Δ*bamE* strain with the plasmid in which expression of the BamE variants was controlled by the BamA promoter utilized as a constitutive promoter. All BamE variants showed similar growth as the WT BamE expressing cell in absence of vancomycin condition (Fig. 3 B). While all mutants of BamE showed growth defect in presence of vancomycin, Y28G showed significant growth defect as much as empty vector (Vector) indicating that Y28 was important for retaining the OM integrity (Fig. 3 C).

**Figure 2.**
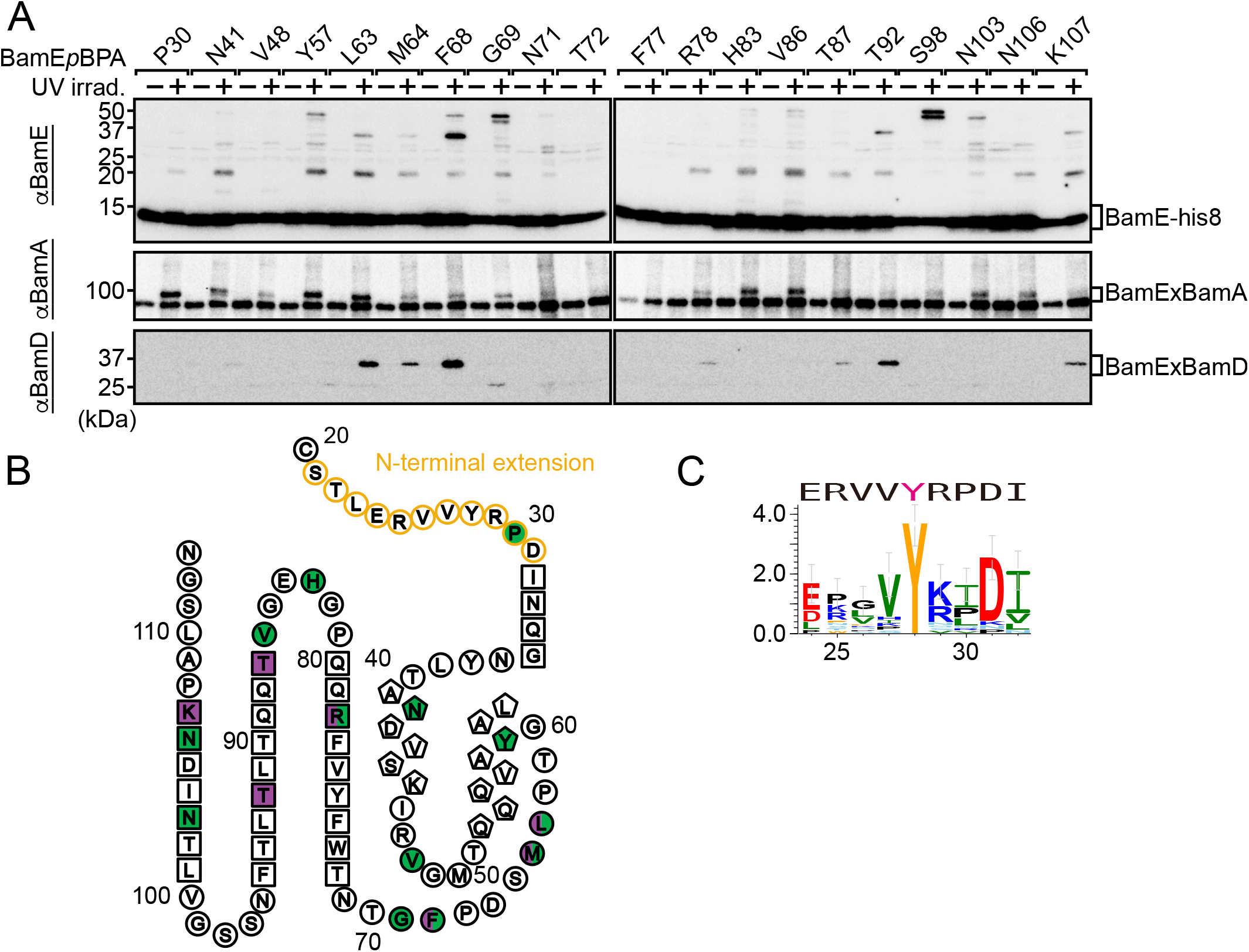
T BamE interacts with BamA by N-terminal extension. (A) *In situ* photo-crosslinking analysis of *p*BPA at the indicated positions of BamE. The cross-linked products were detected with anti-BamE antisera (top), anti-BamA antisera (middle), or anti-BamD antisera (bottom). Immunoblots are representative of at least two biological replicates. (B) Interaction map of BamA on its schematic secondary structure, showing BamE-BamA and BamE-BamD cross-linked sites in green and purple, respectively. Amino acids in the N-terminal extension region are indicated in orange. Circles, pentagons, and squares indicate residues in the loop, α helix, or β strand, respectively. (C) Sequence logo illustrates the conservation of residues 24-32 of BamE; Aromatic (orange), hydrophobic (green), basic (blue), acidic (red), polar (sky blue), and proline and glycine (black). The BamE sequence of E. coli is shown above.

**Figure 3.**
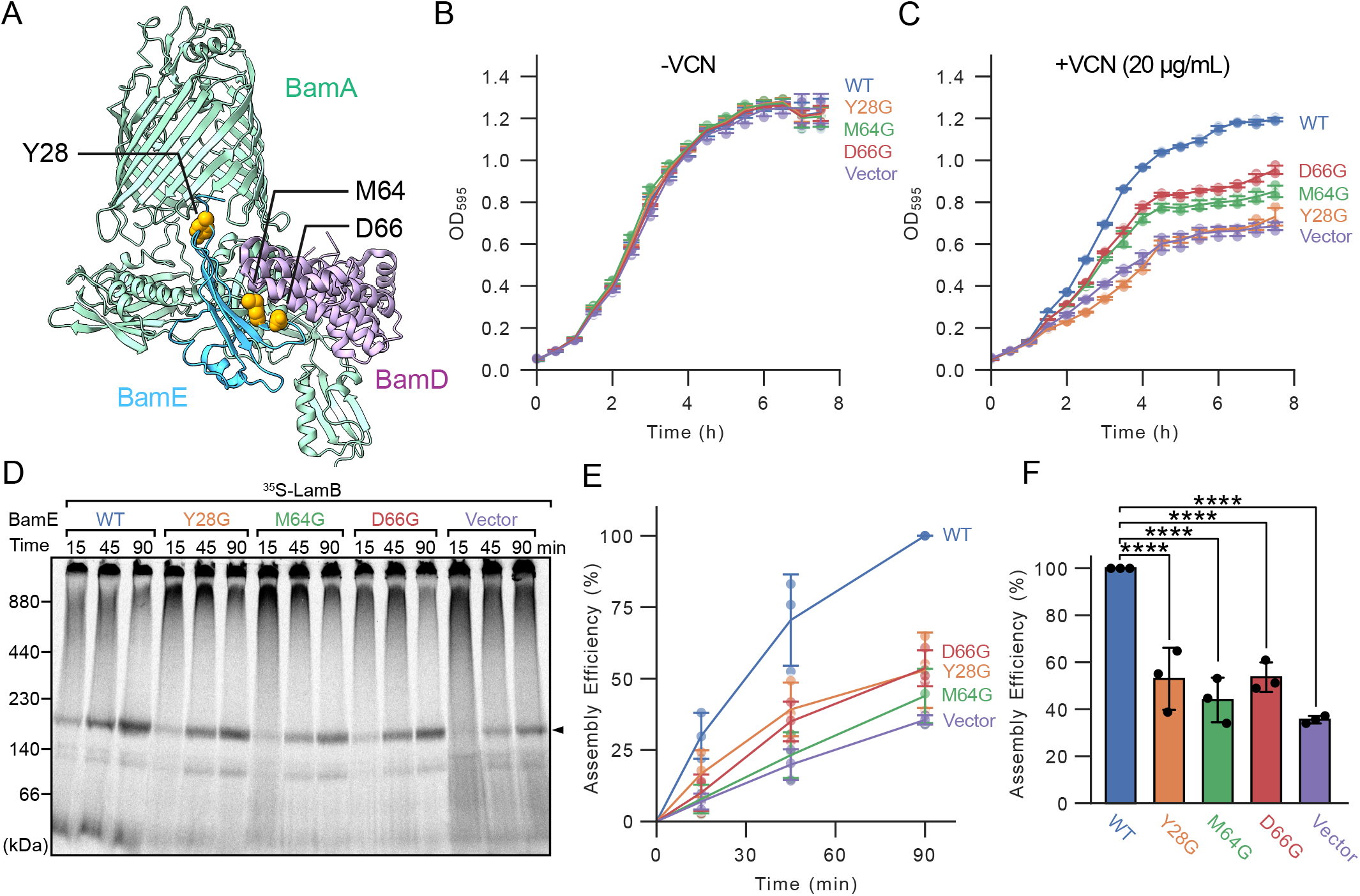
BamE variants affect OM integrity and OMPs assembly efficiency. (A) Mutation sites are indicated in orange on the structure of the BAM complex (5AYW). BamB and BamC were not displayed. (B and C) Δ*bamE* expressing indicated BamE variants were grown in the absence (B) or presence (C) of 20 μg/mL vancomycin (VCN). OD_595_ was measured every 30 minutes. Error bars represent standard deviation from three biological replicates. (D) EMM assembly assay of LamB with BamE variants. Arrowhead indicates assembled trimeric-LamB. All experiments were performed with three technical replicates. (E) Densitometry analysis of D. Amount of trimeric-LamB at 90 minutes in WT set to 100%. Error bars represent standard deviation from three technical replicates. (F) Statistical analysis of assembly efficiency of trimeric-LamB at 90 minutes in D. Error bars represent standard deviation from three technical replicates. Statistical significance was indicated by following signals: ns (not significant), p ≤ 0.10; *, p ≤ 0.05; **, p ≤ 0.01; ***, p ≤ 0.001; ****. Exact values, WT vs Y28G: p=4.74e-5, WT vs M64G: p=2.96e-6, WT vs D66G: 6.37e-5, WT vs Vec: p=4.42e-7 (Dunnett’s multiple comparison test).

To confirm that the decreasing OM integrity of Y28G was due to an assembly defect, we directly analyzed the assembly ability of each bamE mutant by the in vitro reconstitution assay. To reconstitute the assembly reaction, we employed a native membrane fraction, termed *E. coli* Microsomal Membrane (EMM), containing an intact BAM complex (Aoki et al., 2024; Gunasinghe et al., 2018). We isolated EMM from the strains used in the growth curve experiment as in Fig. 3 B and C. The steady-state level of the subunits of the BAM complex in EMM was not different in each variants of BamE when normalized by the level of BamA in EMM (Fig. S5 A and B). Incubation of the radiolabeled substrate with EMM enables the reconstitution of the assembly reaction mediated by the intact BAM complex present in the EMM. For model substrate proteins, we utilized LamB, the 18-stranded OMP and OmpF, the 16-stranded OMP. Our previous study demonstrated that these two substrates significantly required BamE for their assembly (Thewasano et al., 2023). Correctly assembled OmpF and Lamb form a homotrimer which is observed at approximately 160 kDa by Blue Native-PAGE analysis following n-Dodecyl-β-D-maltoside (DDM) solubilization. Densitometry comparison of the 160 kDa LamB homotrimer in each EMM showed that the assembly efficiency in all bamE mutants was reduced to approximately 50% of WT, while the empty vector was reduced to 35% of WT (Fig. 3 D-F). A comparable but somewhat less pronounced effect was observed when OmpF was used as a substrate (Fig. S5 C-E). These results suggest that the highly conserved Y28 in the extension region of BamE is important for substrate assembly.

### The N-terminal extension region modulates the interaction between BamD and the β-barrel domain of BamA

To understand the function of the N-terminal extension region of BamE in substrate assembly, we analyzed the impact of the Y28G mutation on BamA-BamD binding. First, we assessed the stability of the BAM complex using BN-PAGE following DDM solubilization. The fully assembled BAM complex, including all subunits, is typically observed at around 230 kDa (Webb et al., 2012b). When the BamA-BamD interaction is disrupted, the BamA-BamD unit appears as a smaller complex of approximately 140 kDa (Thewasano et al., 2023). In this study, the amount of BamE expressed from the plasmid was insufficient, affecting the stoichiometry of the BAM complex, which resulted in the presence of BamA at both 230 kDa and 140 kDa, even in samples containing WT BamE (Fig. 4A). However, since the 230 kDa complex completely disappeared in the empty vector sample, we conclude that BamE stabilizes the BamA-BamD interaction. Thus, the presence or absence of the 230 kDa complex can be used to assess the role of BamE in stabilizing the BamA-BamD binding. Loss of 230 kDa complex in M64G and D66G indicating that the BamA binding region of the periplasmic domain of BamE contributes to stabilizing the BamA-BamD binding, consistent with previous reports (Kumar and Konovalova, 2023). Interestingly, the Y28G mutant formed a 230 kDa complex, suggesting that the impact of the N-terminal extension region of BamE on substrate assembly differs from that of the periplasmic domain, even though all mutants exhibited similar defects in substrate assembly in the EMM assembly assay.

**Figure 4.**
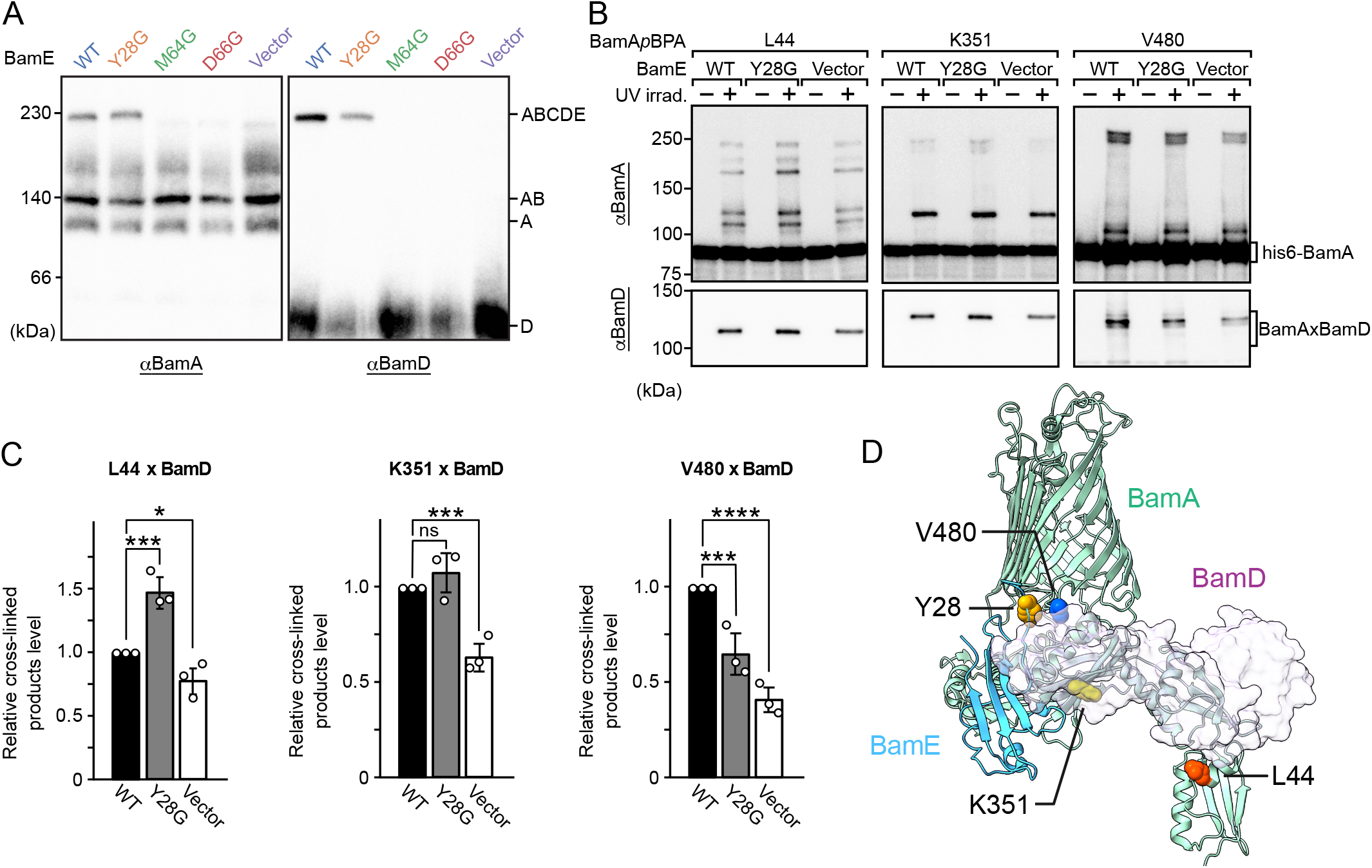
Y28 of BamE fine-tunes proper BamA-BamD interactions. (A) EMM proteins were analyzed by Blue native (BN)-PAGE and immunoblotting using anti-BamA (left) and anti-BamD (right) antisera, respectively. The subunits in each band are shown alphabetically on the right. (B) *In situ* photo-crosslinking analysis of *p*BPA at the indicated positions of BamA in ΔbamE expressing BamE variants. BamA-BamD cross-linked products were compared with WT, Y28G, and empty vector. Cross-linked products were detected with anti-BamA antisera (top) or anti-BamD antisera (bottom). All experiments were performed with three technical replicates. (C) Statistical analysis of BamA-BamD cross-linked products detected by anti-BamD antiserum. Error bars represent standard deviation from three technical replicates. Statistical significance was indicated by signals as Fig 3 F. Exact values, WT vs Y28G at 44 amb: p=2.22e-3, WT vs Vec at 44 amb: p=0.0893, WT vs Y28G at 351 amb: p=0.429, WT vs Vec at 351 amb: p=4.50e-3, WT vs Y28G at 480 amb: p=5.64e-3, WT vs Vec at 480 amb: p=3.37e-4 (Dunnett’s multiple comparison test). (D) Mapping of BamA-BamD cross-linked residues and Y28 of BamE onto the structure of BAM complex (5AYW). BamA and BamD are shown by ribbon and surface model, respectively. Colored residues indicate changes in BamA-BamD cross-linked products in Y28G strain compared to WT. Blue: decreased, Red: increased, Yellow: unchanged. Y28 residue of BamE was indicated in orange.

To further understand the molecular mechanisms of the stimulation of assembly by the N-terminal extension region of BamE, we analyzed the impact of Y28G mutant on local binding of BamA and BamD. To assess the proximity of BamD to BamA, we again employed *in situ* photo cross-linking. We selected three representative sites in BamA, where cross-link products were prominently formed with BamD, in our BamA interaction mapping in Fig. 1A-C and S2. The first site, L44, is located in POTRA 1 and positioned near the N-terminal region of BamD based on the structural data. The second site, K351 in POTRA 5, is adjacent to E373, a key residue involved in the interaction with BamD. The third site, V480 in turn 2 of the β-barrel domain, lies close to Y28 of BamE, indicating a potential interaction interface. Cross-linking experiments at these three positions on BamA were performed using three BamE variants: WT, Y28G, and the null mutant (vector). The levels of each cross-linked product were then compared to evaluate the binding state between BamA and BamD. In the null mutant, the cross-linked product decreased at all positions, with the most pronounced reduction observed in the β-barrel region (V480), consistent with the results shown in Fig. 1 A (Fig. 4 B-D). In the strain expressing the Y28G mutant, the cross-linked product at K351 remained unchanged, while it increased by 50% at L44 and decreased by 35% at V480 (Fig. 4 B-D). These results suggest that the Y28G mutation disrupts the stability of the interaction between BamD and the β-barrel region of BamA, while preserving the interaction between BamD and POTRA 5 of BamA. Additionally, the loss of proximity between BamD and the β-barrel region appears to relatively correlate with an increase in BamD’s proximity to POTRA 1. In summary, our findings indicate that BamE plays two critical roles in facilitating efficient assembly: (1) stabilizing the interaction between BamA and BamD in the periplasmic region, and (2) positioning BamD near the β-barrel domain of BamA through its N-terminal extension region.

### BamD contacts the β-barrel domain of BamA in response to substrate assembly

Our findings thus far demonstrate that the N-terminal extension of BamE is crucial for bringing BamD into proximity with the β-barrel domain of BamA, facilitating efficient assembly. We next investigated whether this proximity correlates with the substrate assembly process. To test this, local proximity was evaluated using *in situ* photo-crosslinking under conditions where assembly was induced by overexpression of a substrate protein (Fig. 5 A). For this purpose, we used OmpC—one of the most abundant major porins—as a model substrate. Additionally, to assess cross-linking under intermediate assembly conditions, we employed both the wild-type (WT) substrate and the FY mutant (F280A and Y286A), a mutant of the internal signal sequence that we previously reported (Germany et al., 2024). The FY mutant stalls in the periplasmic region, lacking the ability to complete pre-folding at BamD. The POTRA 5 and BamD change conformation when they recognize the β-signal of the substrate together (Sinnige et al., 2015). At the POTRA 5 (K351), the crosslinked product with BamD was reduced to half of the empty vector control in overexpression of either substrate. This changing indicates that overexpression of substrate positively produced a substrate-binding state, which altered the proximity state of BamD-BamA. Because FY mutant has a similar affinity to POTRA5 and BamD to WT by containing intact the β-signal, the reduction rate of cross-linked products of POTRA5 and BamD was comparable for both substrates (Fig. 5 A and B). On the other hand, the proximity of the β-barrel domain to BamD changed greatly depending on WT or FY. Overexpression of WT OmpC substrates led to a significant increase in the crosslinking of the β-barrel domain of BamA (V480) to BamD (four-folds) and BamE (12-folds) (Fig. 5). In strains expressing the FY mutant, similar changes were observed, though to a lesser extent compared to the WT (to BamD; two-folds, to BamE; 2.5-folds). In the FY mutant, the substrate stalls at the BamD region and does not trigger the final insertion step, suggesting that the cross-link product between the β-barrel domain and BamD was not fully formed in FY as compared to WT. These results suggest that the proximity of BamD to the β-barrel domain of BamA is dynamically changed in response to the substrate assembly process, with the N-terminal extension region of BamE regulating this proximity to facilitate efficient substrate integration.

**Figure 5.**
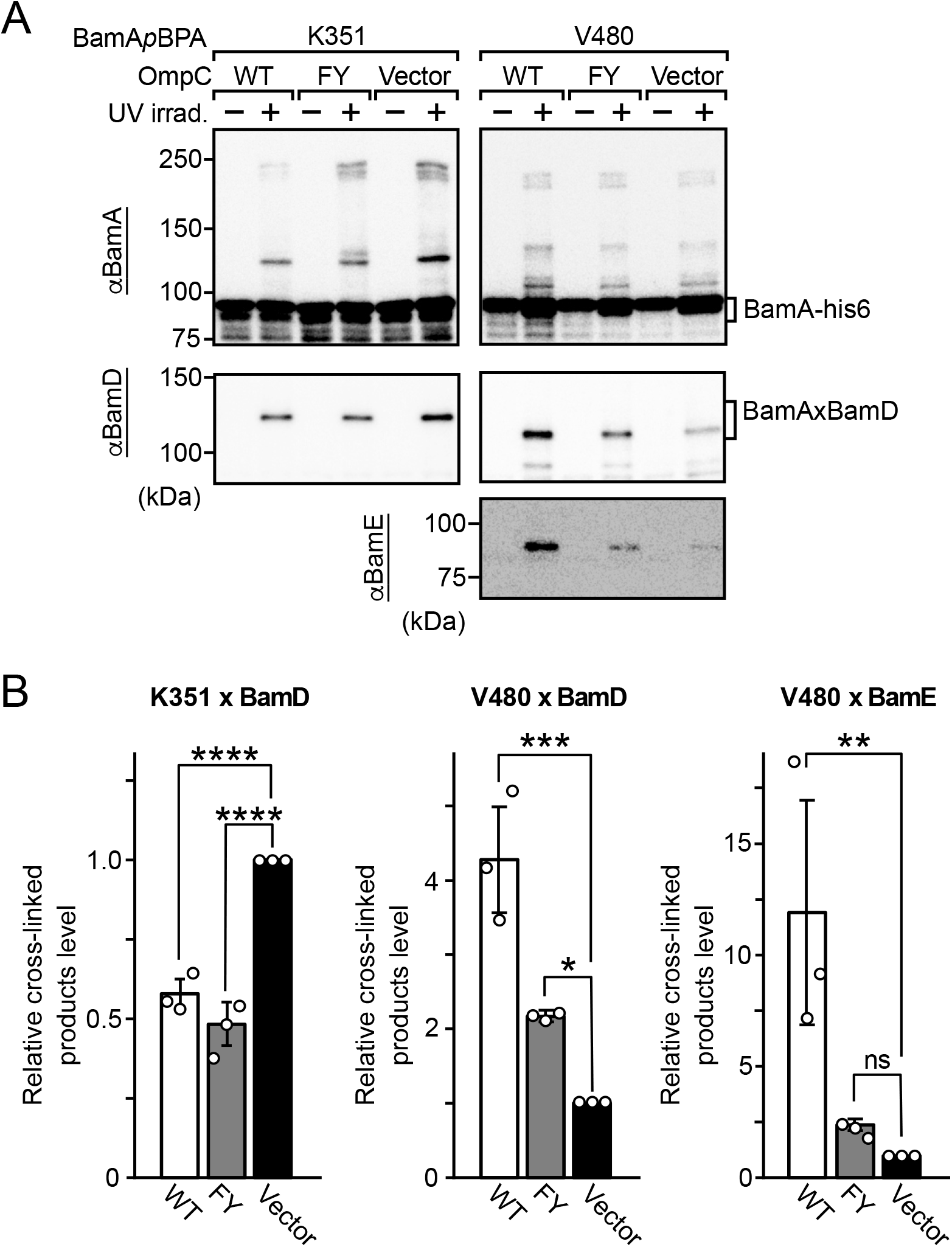
Changing the interaction of BamA and BamD depending on overexpression of various substrates. (A) *In situ* photo-crosslinking analysis of *p*BPA at the indicated positions of BamA in BL21 overexpressing WT or FY mutant of OmpC. Cross-linked products were detected with anti-BamA antisera (top), anti-BamD antisera (middle), and anti-BamE-antisera (bottom). All experiments are performed with three technical replicates. (B) Statistical analysis of BamA-BamD or BamA-BamE cross-linked products detected by anti-BamD or anti-BamE antiserums. Error bars represent standard deviation from three technical replicates. Statistical significance was indicated by signals as Fig. 3 F. Exact values, Vec vs WT at 351 amb: p=2.26e-4, Vec vs FY at 351 amb: p=7.07e-5, Vec vs WT at 480 amb: p=3.86e-4, Vec vs FY at 480 amb: p=0.0524, Vec vs WT at 480 amb (BamE): p=0.0168, Vec vs FY at 480 amb (BamE): p=0.853 (Dunnett’s multiple comparison test).

## Discussion

The main components of the BAM complex are two widely conserved essential subunits, BamA and BamD (Anwari et al., 2012; Webb et al., 2012a). BamD is responsible for pre-folding substrate recognition, while BamA assembles substrates through a β-barrel domain (Germany et al., 2024; Noinaj et al., 2014; Tomasek et al., 2020). It has been suggested that the binding stability between BamA and BamD relies on the presence of BamE (Bakelar et al., 2016; Gu et al., 2016; Iadanza et al., 2016). However, some studies have predicted that the interaction between BamA and BamD requires more precise regulation, as mutations that disrupt BamA-BamD binding cannot be fully restored by compensatory mutations (McCabe et al., 2017a; Ricci et al., 2012; Rigel et al., 2012).

In this study, we identified an additional role for BamE, the regulation of the interaction between BamD and the β-barrel domain of BamA. BamE performs this by modulating the interaction between its N-terminal extension region and turn 2 of BamA. A mutation at the highly conserved residue (Y28G) in BamE reduced OM integrity and impaired substrate assembly *in vitro* without compromising BamA-BamD stability (Fig. 2 C, 3 B-F, and 4 A). Local interaction analysis using site-specific photo-cross-linking revealed that introducing *p*BPA into V480 (turn 2 of BamA) or P30 (within the N-terminal extension of BamE) resulted in a cross-linked product (Fig. 1 A, S1, S3, 2 A and B). This cross-linking product aligns with multiple structural models, indicating that the N-terminal extension of BamE coordinates the interaction between BamD and the β-barrel domain of BamA to facilitate OMPs assembly (Bakelar et al., 2016; Doyle et al., 2022; Fenn et al., 2024; Gu et al., 2016; Han et al., 2016; Haysom et al., 2023; Iadanza et al., 2016; Kaur et al., 2021; Miller et al., 2022; Seyfert et al., 2023; Shen et al., 2023; White et al., 2021; Wu et al., 2021; Xiao et al., 2021).

Since the method to capture the intermediate of substrate transfer from BamD to the β-barrel domain of BamA has not been elucidated in detail yet, structural analysis of this step’s intermediate form has not been dissolved. Thus, the extent to which BamD and the β-barrel domain of BamA must closely cooperate together is currently unknown. The stabilization of BamD binding to the β-barrel domain of BamA by BamE provides an increase in assembly efficiency. This proximity was experimentally increased by inducing the assembly state through substrate overexpression (Fig. 5 A-C). Our study demonstrates that binding of BamD and the β-barrel domain of BamA (at V480) was highest in overexpression of WT OmpC, followed by internal signal mutant (FY mutant) of OmpC that did not complete the assembly, and least in conditions in which the substrate was not expressed. On the other hand, the results of L44 which is the furthest away from the β-barrel domain, were vice versa. Taken together, these results suggest the following conformational changes during assembly (Fig. 6 D): (i) In the resting state, BamD is positioned near the POTRA 1 domain of BamA to capture incoming substrates. (ii) Once the substrate is captured, BamD shifts toward the β-barrel domain of BamA. (iii) BamE stabilizes the interaction between BamD and the β-barrel domain during substrate transfer to the lateral gate of BamA.

**Figure 6.**
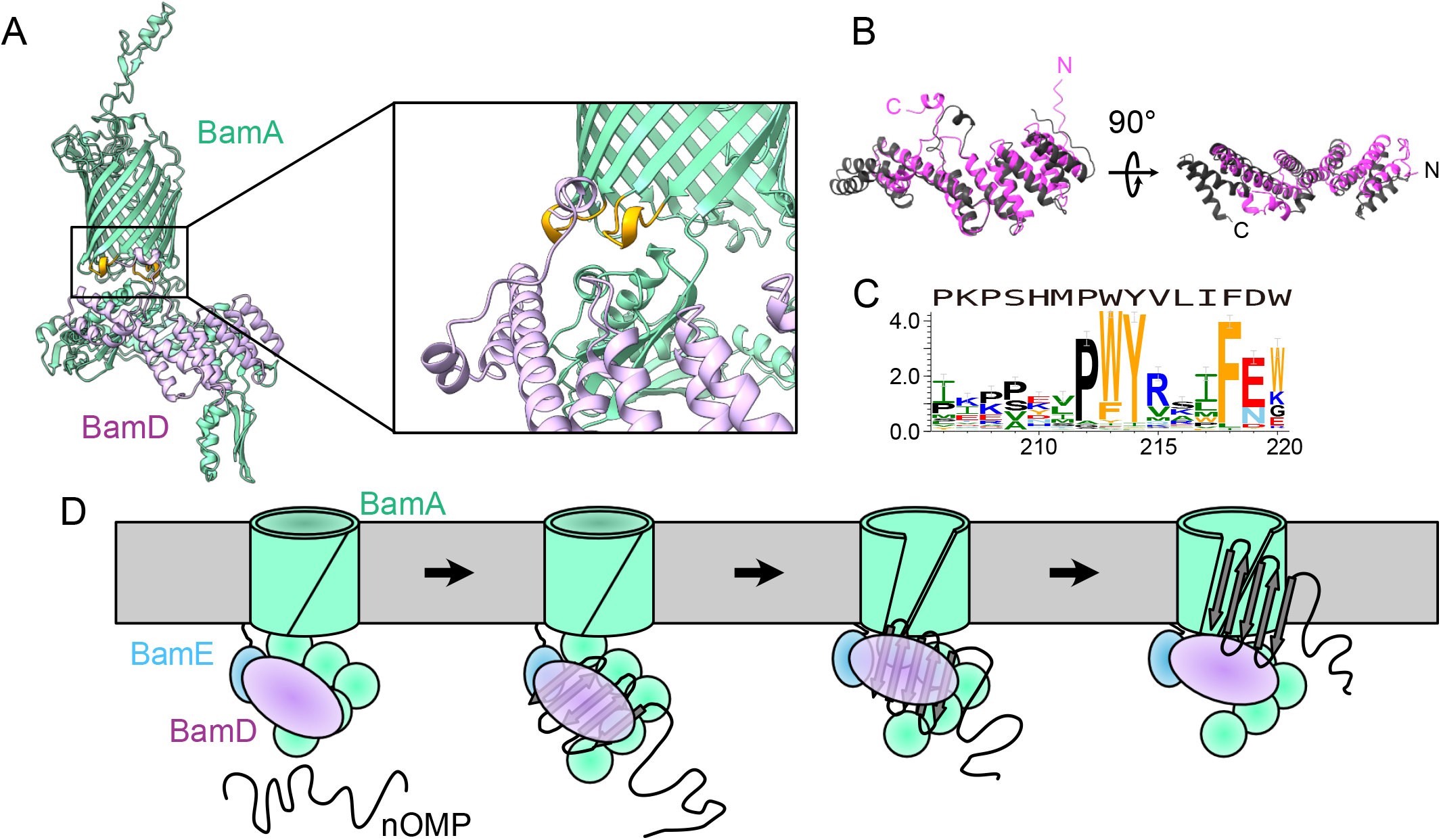
Model diagram of the movement from the BamD recognizing the OMP to passing it on to the BamA. (A) The predicted structure of the BAM complex in H. pylori using AlphaFold3. turn 1 and turn 2 of BamA are shown in orange. (B) Comparison of BamD of *E. coli* (dark gray) and that of H. pylori (magenta) (C) Sequence logo shows the conservation of sequence for residues 206-220 of BamD in epsilon-proteobacteria. Amino acids are colored as described in Fig. 3 C. The BamD sequence of *H. pylori* is shown above the sequence logo. (D) Schematic model of BamD movement during OMPs assembly. nOMP means nascent OMP.

BamE is highly conserved among alpha, beta, and gamma-proteobacteria but is absent in delta- and epsilon-proteobacteria (Anwari et al., 2012). This raises the question: how is binding stabilized and fine-tuned during BamD-to-BamA substrate transfer in bacteria lacking BamE? To address this, we selected Helicobacter pylori as a model bacterium of the epsilon-proteobacteria and predicted the structure of the H. pylori BAM complex composed of BamA and BamD, using AlphaFold3 (Abramson et al., 2024). Interestingly, while both *E. coli* and H. pylori BamD have in common the fold of multiple tetratricopeptide repeat (TPR) motifs, H. pylori BamD was unstructured in the region corresponding to the fifth TPR motif of *E. coli* and interacted with turn 1 and turn 2 of BamA (Fig. 6 A and B). Furthermore, the predicted structures of the BAM complexes of 11 epsilon-proteobacteria indicate that the C-terminal unstructured region of BamD is in close proximity to the Turn region of BamA (Fig. S6). This C-terminal unstructured region contains highly conserved aromatic residues, which may serve a similar function to the N-terminal extension of BamE (Fig. 6 C). Previous analyses have shown that bacteria lacking BamB or BamC possess RmpM or BamF as alternatives to each other, but no alternative subunit to BamE has been found in bacteria lacking BamE (Anwari et al., 2012; Volokhina et al., 2009). Taken together with our findings, the C-terminal unstructured region of BamD in epsilon-proteobacteria may compensate for the absence of BamE, providing the required stabilization and fine-tuning for efficient substrate transport and assembly.

## Materials and Methods

### Bacterial strains and growth conditions

Bacterial strains used in this study were listed in S1 Table. *E. coli* cells were incubated at 37°C in L medium (10 g/L tryptone, 5g/L yeast extract, and 10 g/L NaCl) with appropriate antibiotics (100 μg/mL ampicillin, 30 μg/mL kanamycin and 50 μg/mL chloramphenicol, 50 μg/mL spectinomycin) for selecting transformants.

### Plasmids and primers

Plasmids and primers used in this study were listed in S2 and S3 Table, respectively. Point mutation was introduced into pTnT-bamEhis8 via Quick change method (Imai et al., 1991). These point mutations were confirmed by sequencing. The spectinomycin resistant plasmids to express the BamA (X amb) derivatives were listed in S2 Table, and constructed as follows. To swap from ampicillin marker to spectinomycin marker, we amplified spectinomycin resistant region using a pair of pET-Spe^R^1 and pET-Spe^R^2, and the plasmids except for ampicillin resistant region were amplified by a pair of primer pET-Spe^R^3 and pET-Spe^R^4. Both DNA fragments were combined by SLiCE method (Motohashi, 2015). The sequence of spectinomycin resistant gene was shown in S4 Table.

### Isolation of EMM

EMM was prepared as described previously (Aoki et al., 2024; Gunasinghe et al., 2018; Thewasano et al., 2023). In brief, *E. coli* cells were incubated at 37°C to an OD_595_ of ∼ 1.0. Cells were harvested by centrifugation (4500 rpm, 7 min, 4°C). The pellets were resuspended in sonication buffer (50 mM Tris-HCl pH 7.5, 150 mM NaCl, and 5 mM EDTA) and sonicated on ice. Treated cell mixtures were centrifuged (3000 rpm, 5 min, 4°C), and the supernatant was pelleted by centrifuge (11,000 rpm, 10 min, 4°C). The pellet was dissolved in SEM buffer (250 mM Sucrose, 10 mM MOPS-KOH pH 7.2, and 1 mM EDTA) and flash-frozen in liquid nitrogen, and stored at - 80°C. Protein concentrations of EMM were determined by Absorbance 280 nm measurements of 100-fold diluted in 0.6% SDS (Absorbance 280 nm = 0.21, set to 10 mg/ml).

### Immunoblotting

Total proteins were separated by SDS-PAGE, and electro-transferred onto a polyvinylidene difluoride membrane. The membrane was treated with 2% w/v skim milk in TBST (20 mM Tris-HCl pH 8.0, 150 mM NaCl, 0.05% v/v Tween 20), and then inundated with rabbit anti-BamA (1/10000 dilution), rabbit anti-BamB (1/20000 dilution), rabbit anti-BamC (1/10000 dilution), rabbit anti-BamD (1/10000 dilution), mouse anti-BamD (1/1000 dilution), or rabbit anti-BamE (1/10000 dilution) in 2% skim milk in TBST. All primary antibodies were gifted from Trevor Lithgow laboratory (Monash University). After washing with TBST twice, the membranes were incubated with a HRP-conjugated goat anti-rabbit IgG (1/20000 dilution) or a HRP-conjugated goat anti-mouse IgG (1/20000 dilution) in 2% w/v skim milk in TBST. Proteins were visualized with Chemi-Lumi One usingWSE-6270LuminoGraph Ⅱ EM.

### Steady-state protein level and protein complex analysis

To check steady-state levels of BAM complex subunits, EMM pellets were resuspended in SDS-PAGE sample buffer (125 mM Tris-HCl, pH 6.8, 2 mM EDTA, 2% w/v SDS, 1% w/v sucrose, 0.03% w/v bromophenol blue, 1% v/v β-mercaptoethanol). The samples were boiled at 98°C for 5 min, centrifuged at 12,000 rpm for 5 min at 25°C, then subjected to SDS-PAGE. Afterward, the levels of the BAM complex subunits were analyzed by immunoblotting with anti-BamA, anti-BamB, anti-BamC, anti-BamD, and anti-BamE antisera.

To check whether BamE variants form the BAM complex, EMM pellets were solubilized with 1.5% DDM containing BN-PAGE lysis buffer (25 mM imidazole-HCl, pH 7.0, 50 mM NaCl, 50 mM 6-aminohexanoic acid, 1 mM EDTA, 7.5% w/v glycerol) on ice for 20 min. Solubilized proteins were clarified by centrifugation (15,000 rpm, 10 min, 4°C), mixing with 20x BN-PAGE sample buffer (4.0% w/v CBB G-250, 100 mM 6-aminohexanoic acid), and subjected to BN-PAGE. After BN-PAGE electrophoresis, the gel was incubated in denaturing buffer (200 mM Tris-HCl pH 7.2, 4% w/v SDS, 0.5% v/v β-mercaptoethanol), and then the proteins were transferred to a PVDF membrane. PVDF membrane transferred proteins were washed with 100% methanol and immunoblotted with anti-BamA and anti-BamD antisera.

### BamA interaction mapping

BamA shut-off cells carrying pSup-BpaRS-6TRN and pTnT-H6A2bamA (X amb) were incubated at 37°C in L medium containing 0.02% w/v arabinose. The overnight culture was transferred into X-link medium (10g/L tryptone, 1g/L yeast extract, 10 g/L NaCl, and 1 mM *p*BPA) containing 0.2% w/v glucose and the cells were grown at 37°C for 5 h in the dark.

After incubation in X-link medium, the half volume of the cultures was UV irradiated. The other half was kept on ice as non-UV-irradiated samples. After UV-irradiation, the cultures were centrifuged (4500 rpm, 7 min, 4°C). The pellets were disrupted by sonication in 1% SDS buffer (50 mM Tris-HCl pH 8.0, 150 mM NaCl, 1% SDS). His-tagged proteins which contain *p*BPA, and its cross-linked products were purified with Ni-NTA agarose. Purified samples were mixed with 3 x SDS sample buffer. These samples were subjected to SDS-PAGE and assessed by immunoblotting with anti-BamA, anti-BamD, or anti-BamE antisera.

### BamE interaction mapping

Δ*bamE* cells carrying pEVOL-BpF and pTnT-bamE-his8 (X amb) were incubated at 37°C in L medium. The overnight culture was transferred into X-link medium containing 0.2% w/v arabinose and the cells were grown at 37°C for 5 h in the dark. After incubation, UV was irradiated and cross-linked products were assessed by immunoblotting as described above.

### BamA-BamD interaction analysis in BL21 vs Δ*bamE*

BL21 and Δ*bamE* cells carrying pEVOL-BpF and pTnT-H6A2bamA (X amb) were incubated at 37°C in L medium. The overnight culture was transferred into X-link medium containing 0.2% w/v arabinose and the cells were grown at 37°C for 5 h in the dark. After incubation, UV was irradiated and cross-linked products were assessed by immunoblotting as described above.

### BamA-BamD interaction analysis in BamE mutant

Δ*bamE* cells carrying pEVOL-BpF, pTnT (Spe^R^)-H6A2bamA (44, 351, or 480 amb), and pTnT-bamE-his8 (WT, Y28G, or empty vector) were incubated at 37°C in L medium. The overnight culture was transferred into X-link medium containing 0.2% w/v arabinose and the cells were incubated at 37°C for 5h in the dark. After incubation, UV was irradiated and cross-linked products were assessed by immunoblotting as described above.

### BamA-BamD interaction analysis with overexpressing OmpC

BL21 cells harboring pEVOL-BpF, pTnT (Spe^R^)-H6A2bamA (44, 351, or 480 amb), and pBAD-FLOmpC (WT, F280A Y286A, or empty vector) were incubated at 37°C in L medium. The overnight culture was transferred into X-link medium for 3.5 h. Next, arabinose was supplemented at a final concentration of 0.2% w/v and the cells were incubated for 1.5 h. After incubation, UV was irradiated and cross-linked products were assessed by immunoblotting as described above.

### EMM assembly assay

Experimental procedures of EMM assembly assay were described previously (Aoki et al., 2024; Gunasinghe et al., 2018; Thewasano et al., 2023). In brief, OmpF and LamB were prepared by in vitro translation in rabbit reticulocyte lysate supplemented with ^35^S-methionine using in vitro transcribed mRNA by SP6-RNA polymerase. 150 μg EMM were resuspended in 200 μl assembly assay buffer (10 mM Mops-KOH pH 7.2, 2.5 mM KH_2_PO_4_, 250 mM sucrose, 15 mM KCl, 5 mM MgCl_2_, 2 mM methionine, 5 mM DTT, 1% w/v BSA, 0.09% v/v Triton X-100) incubated with 20 μl ^35^S-labeled OmpF or LamB per a reaction at 30°C. The assembly reactions were halted at 15, 45, or 90 min by shifting on ice. The harvested EMM by centrifugation (14,000 rpm, 5 min, 4°C) were washed with SEM buffer, and then EMM pellets were solubilized with 1.5% DDM containing BN-PAGE lysis buffer subjected to BN-PAGE as described above. After BN-PAGE electrophoresis, the gel was dried and exposed on IP plates.

### Sequence conservation analysis

For conservation analysis, BLAST was used to obtain homologous sequences of BamE and BamD from swissprot and RefSeq Select proteins, respectively. The hit sequences of BamE in epsilon-proteobacteria were used. We identified 21 and 182 sequences for BamE and BamD, respectively. Multiple sequence alignments were generated by Clustal Omega and then sequence logos were created using WebLogo 3 (Crooks et al., 2004).

### Densitometry and statistical analysis

For the EMM assembly assay and steady-state level protein comparison, three independent experimental datasets were densitometrically measured using CS Analyzer software from Atto. Statistical analyses were performed with Dunnett’s multiple comparison test based on three independent technical replicates to compare more than three groups. Statistical significance was set at p values < 0.10 and denoted with an asterisk. Exact *p* values of each experiment were described in the figure legends.

## Acknowledgments

We thank the members of the Shiota labs, Eriko Aoki at Tottori University, Hotomi Mimuro, and Tomohiro Miyoshi at Oita University for discussion and critical comments on the manuscript. We appreciate the Trevor Lithgow lab for providing us with the BAM accessory protein deletion mutants, and antibodies against subunits of the BAM complex. Radiation experiments were supported by the RI Kiyotake of the Frontier Science Research Center, University of Miyazaki. This work was partly supported by the Research Center for GLOBAL and LOCAL Infectious Diseases, Oita University (2024B06). T.S. was supported by JSPS KAKENHI (22K12672 and 24KK0138), the JST FOREST Program (JPMJFR2064), and Takeda Science Foundation. E. M. G. was supported by JSPS KAKENHI (24K18071).

## Notes

### Competing Interest Statement

The authors have declared no competing interest.

## References

Abramson J, Adler J, Dunger J, Evans R, Green T, Pritzel A, Ronneberger O, Willmore L, Ballard AJ, Bambrick J, Bodenstein SW, Evans DA, Hung C-C, O’Neill M, Reiman D, Tunyasuvunakool K, Wu Z, Žemgulytė A, Arvaniti E, Beattie C, Bertolli O, Bridgland A, Cherepanov A, Congreve M, Cowen-Rivers AI, Cowie A, Figurnov M, Fuchs FB, Gladman H, Jain R, Khan YA, Low CMR, Perlin K, Potapenko A, Savy P, Singh S, Stecula A, Thillaisundaram A, Tong C, Yakneen S, Zhong ED, Zielinski M, Žídek A, Bapst V, Kohli P, Jaderberg M, Hassabis D, Jumper JM. 2024. Accurate structure prediction of biomolecular interactions with AlphaFold 3. Nature 630:493–500. doi:10.1038/s41586-024-07487-w

Anwari K, Webb CT, Poggio S, Perry AJ, Belousoff M, Celik N, Ramm G, Lovering A, Sockett RE, Smit J, Jacobs-Wagner C, Lithgow T. 2012. The evolution of new lipoprotein subunits of the bacterial outer membrane BAM complex. Mol Microbiol 84:832–844. doi:10.1111/j.1365-2958.2012.08059.x

Aoki E, Germany E, Shiota T. 2024. In Vitro Reconstruction of Bacterial β-Barrel Membrane Protein Assembly Using E. coli Microsomal (Mid-Density) Membrane. Methods Mol Biol Clifton NJ 2778:65–81. doi:10.1007/978-1-0716-3734-0_5

Bakelar J, Buchanan SK, Noinaj N. 2016. The structure of the β-barrel assembly machinery complex. Science 351:180–186. doi:10.1126/science.aad3460

Bos MP, Tommassen J. 2004. Biogenesis of the Gram-negative bacterial outer membrane. Curr Opin Microbiol 7:610–616. doi:10.1016/j.mib.2004.10.011

Chen X, Ding Y, Bamert RS, Le Brun AP, Duff AP, Wu C-M, Hsu H-Y, Shiota T, Lithgow T, Shen H-H. 2021. Substrate-dependent arrangements of the subunits of the BAM complex determined by neutron reflectometry. Biochim Biophys Acta BBA - Biomembr 1863:183587. doi:10.1016/j.bbamem.2021.183587

Chin JW, Martin AB, King DS, Wang L, Schultz PG. 2002. Addition of a photocrosslinking amino acid to the genetic code of Escherichiacoli. Proc Natl Acad Sci U S A 99:11020–11024. doi:10.1073/pnas.172226299

Crooks GE, Hon G, Chandonia J-M, Brenner SE. 2004. WebLogo: a sequence logo generator. Genome Res 14:1188–1190. doi:10.1101/gr.849004

Doyle MT, Bernstein HD. 2022. Function of the Omp85 Superfamily of Outer Membrane Protein Assembly Factors and Polypeptide Transporters. Annu Rev Microbiol 76:259–279. doi:10.1146/annurev-micro-033021-023719

Doyle MT, Jimah JR, Dowdy T, Ohlemacher SI, Larion M, Hinshaw JE, Bernstein HD. 2022. Cryo-EM structures reveal multiple stages of bacterial outer membrane protein folding. Cell 185:1143–1156.e13. doi:10.1016/j.cell.2022.02.016

Fenn KL, Horne JE, Crossley JA, Böhringer N, Horne RJ, Schäberle TF, Calabrese AN, Radford SE, Ranson NA. 2024. Outer membrane protein assembly mediated by BAM-SurA complexes. Nat Commun 15:7612. doi:10.1038/s41467-024-51358-x

Germany EM, Thewasano N, Imai K, Maruno Y, Bamert RS, Stubenrauch CJ, Dunstan RA, Ding Y, Nakajima Y, Lai X, Webb CT, Hidaka K, Tan KS, Shen H, Lithgow T, Shiota T. 2024. Dual recognition of multiple signals in bacterial outer membrane proteins enhances assembly and maintains membrane integrity. eLife 12:RP90274. doi:10.7554/eLife.90274

Gu Y, Li H, Dong H, Zeng Y, Zhang Z, Paterson NG, Stansfeld PJ, Wang Z, Zhang Y, Wang W, Dong C. 2016. Structural basis of outer membrane protein insertion by the BAM complex. Nature 531:64–69. doi:10.1038/nature17199

Guest RL, Silhavy TJ. 2023. Cracking outer membrane biogenesis. Biochim Biophys Acta BBA - Mol Cell Res 1870:119405. doi:10.1016/j.bbamcr.2022.119405

Gunasinghe SD, Shiota T, Stubenrauch CJ, Schulze KE, Webb CT, Fulcher AJ, Dunstan RA, Hay ID, Naderer T, Whelan DR, Bell TDM, Elgass KD, Strugnell RA, Lithgow T. 2018. The WD40 Protein BamB Mediates Coupling of BAM Complexes into Assembly Precincts in the Bacterial Outer Membrane. Cell Rep 23:2782–2794. doi:10.1016/j.celrep.2018.04.093

Hagan CL, Wzorek JS, Kahne D. 2015. Inhibition of the β-barrel assembly machine by a peptide that binds BamD. Proc Natl Acad Sci U S A 112:2011–2016. doi:10.1073/pnas.1415955112

Han L, Zheng J, Wang Y, Yang X, Liu Y, Sun C, Cao B, Zhou H, Ni D, Lou J, Zhao Y, Huang Y. 2016. Structure of the BAM complex and its implications for biogenesis of outer-membrane proteins. Nat Struct Mol Biol 23:192–196. doi:10.1038/nsmb.3181

Hart EM, Gupta M, Wühr M, Silhavy TJ. 2020. The gain-of-function allele bamAE470K bypasses the essential requirement for BamD in β-barrel outer membrane protein assembly. Proc Natl Acad Sci U S A 117:18737–18743. doi:10.1073/pnas.2007696117

Hayashi S, Buchanan SK, Botos I. 2024a. The Name Is Barrel, β-Barrel. Methods Mol Biol Clifton NJ 2778:1–30. doi:10.1007/978-1-0716-3734-0_1

Hayashi S, Buchanan SK, Botos I. 2024b. The Name Is Barrel, β-Barrel. Methods Mol Biol Clifton NJ 2778:1–30. doi:10.1007/978-1-0716-3734-0_1

Haysom SF, Machin J, Whitehouse JM, Horne JE, Fenn K, Ma Y, El Mkami H, Böhringer N, Schäberle TF, Ranson NA, Radford SE, Pliotas C. 2023. Darobactin B Stabilises a Lateral-Closed Conformation of the BAM Complex in E. coli Cells. Angew Chem Int Ed Engl 62:e202218783. doi:10.1002/anie.202218783

Heinz E, Lithgow T. 2014. A comprehensive analysis of the Omp85/TpsB protein superfamily structural diversity, taxonomic occurrence, and evolution. Front Microbiol 5:370. doi:10.3389/fmicb.2014.00370

Iadanza MG, Higgins AJ, Schiffrin B, Calabrese AN, Brockwell DJ, Ashcroft AE, Radford SE, Ranson NA. 2016. Lateral opening in the intact β-barrel assembly machinery captured by cryo-EM. Nat Commun 7:12865. doi:10.1038/ncomms12865

Imai Y, Matsushima Y, Sugimura T, Terada M. 1991. A simple and rapid method for generating a deletion by PCR. Nucleic Acids Res 19:2785. doi:10.1093/nar/19.10.2785

Kaur H, Jakob RP, Marzinek JK, Green R, Imai Y, Bolla JR, Agustoni E, Robinson CV, Bond PJ, Lewis K, Maier T, Hiller S. 2021. The antibiotic darobactin mimics a β-strand to inhibit outer membrane insertase. Nature 593:125–129. doi:10.1038/s41586-021-03455-w

Knowles TJ, Browning DF, Jeeves M, Maderbocus R, Rajesh S, Sridhar P, Manoli E, Emery D, Sommer U, Spencer A, Leyton DL, Squire D, Chaudhuri RR, Viant MR, Cunningham AF, Henderson IR, Overduin M. 2011. Structure and function of BamE within the outer membrane and the β-barrel assembly machine. EMBO Rep 12:123–128. doi:10.1038/embor.2010.202

Kumar S, Konovalova A. 2023. BamE directly interacts with BamA and BamD coordinating their functions. Mol Microbiol 120:397–407. doi:10.1111/mmi.15127

Lehr U, Schütz M, Oberhettinger P, Ruiz-Perez F, Donald JW, Palmer T, Linke D, Henderson IR, Autenrieth IB. 2010. C-terminal amino acid residues of the trimeric autotransporter adhesin YadA of Yersinia enterocolitica are decisive for its recognition and assembly by BamA. Mol Microbiol 78:932–946. doi:10.1111/j.1365-2958.2010.07377.x

McCabe AL, Ricci D, Adetunji M, Silhavy TJ. 2017a. Conformational Changes That Coordinate the Activity of BamA and BamD Allowing β-Barrel Assembly. J Bacteriol 199:e00373–17. doi:10.1128/JB.00373-17

McCabe AL, Ricci D, Adetunji M, Silhavy TJ. 2017b. Conformational Changes That Coordinate the Activity of BamA and BamD Allowing β-Barrel Assembly. J Bacteriol 199:e00373–17. doi:10.1128/JB.00373-17

Miller RD, Iinishi A, Modaresi SM, Yoo B-K, Curtis TD, Lariviere PJ, Liang L, Son S, Nicolau S, Bargabos R, Morrissette M, Gates MF, Pitt N, Jakob RP, Rath P, Maier T, Malyutin AG, Kaiser JT, Niles S, Karavas B, Ghiglieri M, Bowman SEJ, Rees DC, Hiller S, Lewis K. 2022. Computational identification of a systemic antibiotic for gram-negative bacteria. Nat Microbiol 7:1661–1672. doi:10.1038/s41564-022-01227-4

Motohashi K. 2015. A simple and efficient seamless DNA cloning method using SLiCE from Escherichia coli laboratory strains and its application to SLiP site-directed mutagenesis. BMC Biotechnol 15:47. doi:10.1186/s12896-015-0162-8

Noinaj N, Kuszak AJ, Balusek C, Gumbart JC, Buchanan SK. 2014. Lateral opening and exit pore formation are required for BamA function. Struct Lond Engl 1993 22:1055–1062. doi:10.1016/j.str.2014.05.008

Noinaj N, Kuszak AJ, Gumbart JC, Lukacik P, Chang H, Easley NC, Lithgow T, Buchanan SK. 2013. Structural insight into the biogenesis of β-barrel membrane proteins. Nature 501:385–390. doi:10.1038/nature12521

Ricci DP, Hagan CL, Kahne D, Silhavy TJ. 2012. Activation of the Escherichia coli β-barrel assembly machine (Bam) is required for essential components to interact properly with substrate. Proc Natl Acad Sci U S A 109:3487–3491. doi:10.1073/pnas.1201362109

Rigel NW, Schwalm J, Ricci DP, Silhavy TJ. 2012. BamE modulates the Escherichia coli beta-barrel assembly machine component BamA. J Bacteriol 194:1002–1008. doi:10.1128/JB.06426-11

Seyfert CE, Porten C, Yuan B, Deckarm S, Panter F, Bader CD, Coetzee J, Deschner F, Tehrani KHME, Higgins PG, Seifert H, Marlovits TC, Herrmann J, Müller R. 2023. Darobactins Exhibiting Superior Antibiotic Activity by Cryo-EM Structure Guided Biosynthetic Engineering. Angew Chem Int Ed Engl 62:e202214094. doi:10.1002/anie.202214094

Shen C, Chang S, Luo Q, Chan KC, Zhang Z, Luo B, Xie T, Lu G, Zhu X, Wei X, Dong C, Zhou R, Zhang X, Tang X, Dong H. 2023. Structural basis of BAM-mediated outer membrane β-barrel protein assembly. Nature 617:185–193. doi:10.1038/s41586-023-05988-8

Sinnige T, Weingarth M, Daniëls M, Boelens R, Bonvin AMJJ, Houben K, Baldus M. 2015. Conformational Plasticity of the POTRA 5 Domain in the Outer Membrane Protein Assembly Factor BamA. Struct Lond Engl 1993 23:1317–1324. doi:10.1016/j.str.2015.04.014

Sklar JG, Wu T, Gronenberg LS, Malinverni JC, Kahne D, Silhavy TJ. 2007. Lipoprotein SmpA is a component of the YaeT complex that assembles outer membrane proteins in Escherichia coli. Proc Natl Acad Sci U S A 104:6400–6405. doi:10.1073/pnas.0701579104

Thewasano N, Germany EM, Maruno Y, Nakajima Y, Shiota T. 2023. Categorization of Escherichia coli outer membrane proteins by dependence on accessory proteins of the β-barrel assembly machinery complex. J Biol Chem 299:104821. doi:10.1016/j.jbc.2023.104821

Tomasek D, Kahne D. 2021. The assembly of β-barrel outer membrane proteins. Curr Opin Microbiol 60:16–23. doi:10.1016/j.mib.2021.01.009

Tomasek D, Rawson S, Lee J, Wzorek JS, Harrison SC, Li Z, Kahne D. 2020. Structure of a nascent membrane protein as it folds on the BAM complex. Nature 583:473–478. doi:10.1038/s41586-020-2370-1

Volokhina EB, Beckers F, Tommassen J, Bos MP. 2009. The beta-barrel outer membrane protein assembly complex of Neisseria meningitidis. J Bacteriol 191:7074–7085. doi:10.1128/JB.00737-09

Voulhoux R, Bos MP, Geurtsen J, Mols M, Tommassen J. 2003. Role of a highly conserved bacterial protein in outer membrane protein assembly. Science 299:262–265. doi:10.1126/science.1078973

Webb CT, Heinz E, Lithgow T. 2012a. Evolution of the β-barrel assembly machinery. Trends Microbiol 20:612–620. doi:10.1016/j.tim.2012.08.006

Webb CT, Selkrig J, Perry AJ, Noinaj N, Buchanan SK, Lithgow T. 2012b. Dynamic association of BAM complex modules includes surface exposure of the lipoprotein BamC. J Mol Biol 422:545– 555. doi:10.1016/j.jmb.2012.05.035

White P, Haysom SF, Iadanza MG, Higgins AJ, Machin JM, Whitehouse JM, Horne JE, Schiffrin B, Carpenter-Platt C, Calabrese AN, Storek KM, Rutherford ST, Brockwell DJ, Ranson NA, Radford SE. 2021. The role of membrane destabilisation and protein dynamics in BAM catalysed OMP folding. Nat Commun 12:4174. doi:10.1038/s41467-021-24432-x

Wu R, Bakelar JW, Lundquist K, Zhang Z, Kuo KM, Ryoo D, Pang YT, Sun C, White T, Klose T, Jiang W, Gumbart JC, Noinaj N. 2021. Plasticity within the barrel domain of BamA mediates a hybrid-barrel mechanism by BAM. Nat Commun 12:7131. doi:10.1038/s41467-021-27449-4

Xiao L, Han L, Li B, Zhang M, Zhou H, Luo Q, Zhang X, Huang Y. 2021. Structures of the β-barrel assembly machine recognizing outer membrane protein substrates. FASEB J Off Publ Fed Am Soc Exp Biol 35:e21207. doi:10.1096/fj.202001443RR

