## supplemental figure, tables for "The N-terminal extension region of BamE coordinates interaction between the β-barrel domain of BamA and BamD for efficient assembly"

*Takuya Shiota

**Author Contributions:** Y. Maruno and T. S. designed research. Y. Maruno, E. M. G., and T. S. wrote the manuscript. Y. Maruno and T. S. performed most of the key experiments and analyzed data. Y. Maruno, Y. N., R. O., and Y. Masumura constructed plasmid and produced *E. coli* strains. R. O. and Y. Masumura performed interaction mapping.

**Competing Interest Statement:** The authors declare no competing interests.

**Classification:** Biological Science, Biochemistry

**Keywords:** β-barrel assembly machinery, outer membrane proteins, Gram-negative bacteria, protein folding, membrane protein insertion

**This PDF file includes:**

Figures S1 to S6
Tables S1 to S4
SI References

### Supporting Figures

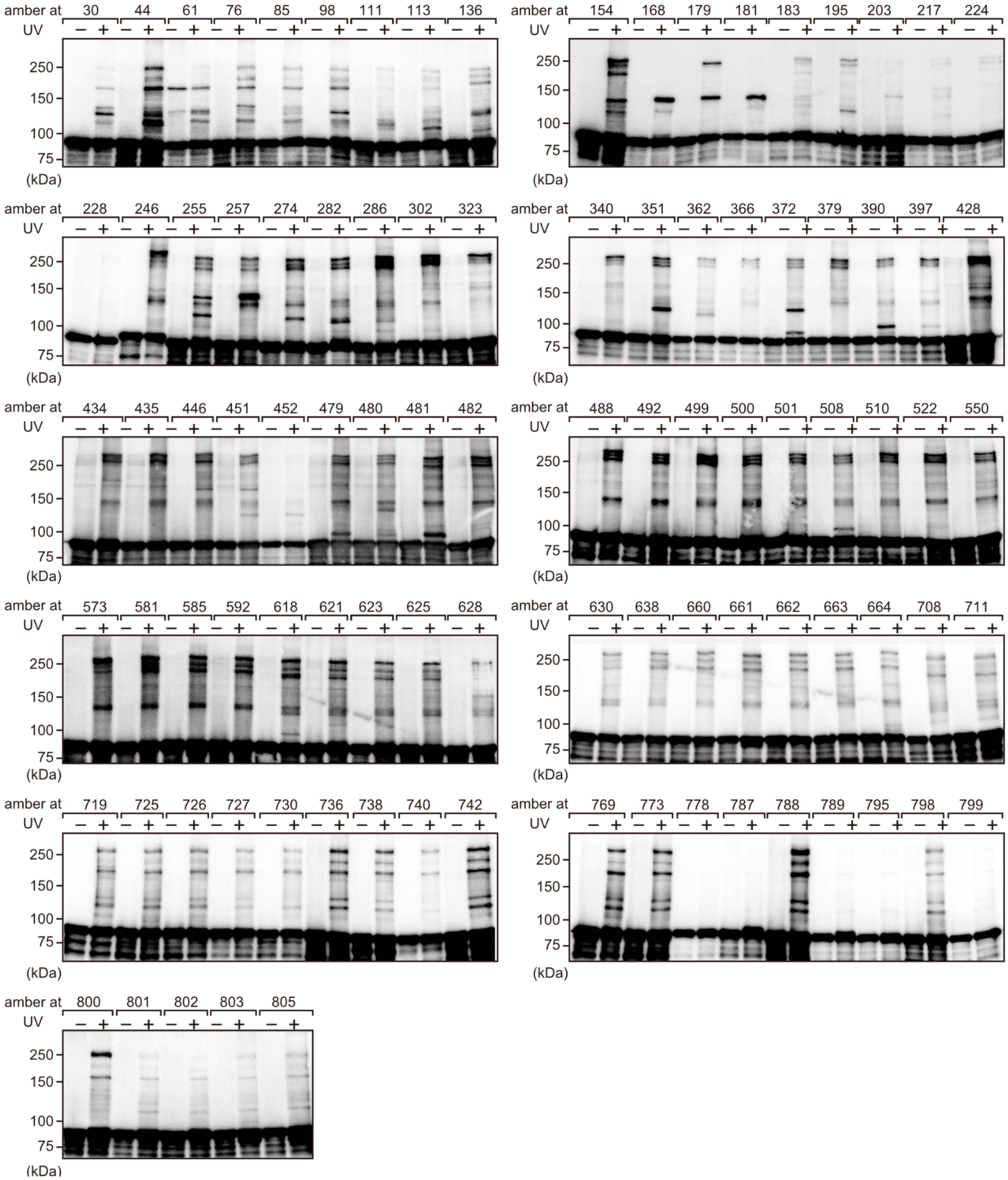


**Fig. S1.** Interaction mapping by *in situ* photo-crosslinking. *p*BPA was introduced at indicated positions of BamA. Hexahistidine-tagged BamA and its cross-linked products were purified with Ni-NTA, and then analyzed by SDS-PAGE and immunoblotting using anti-BamA antisera.

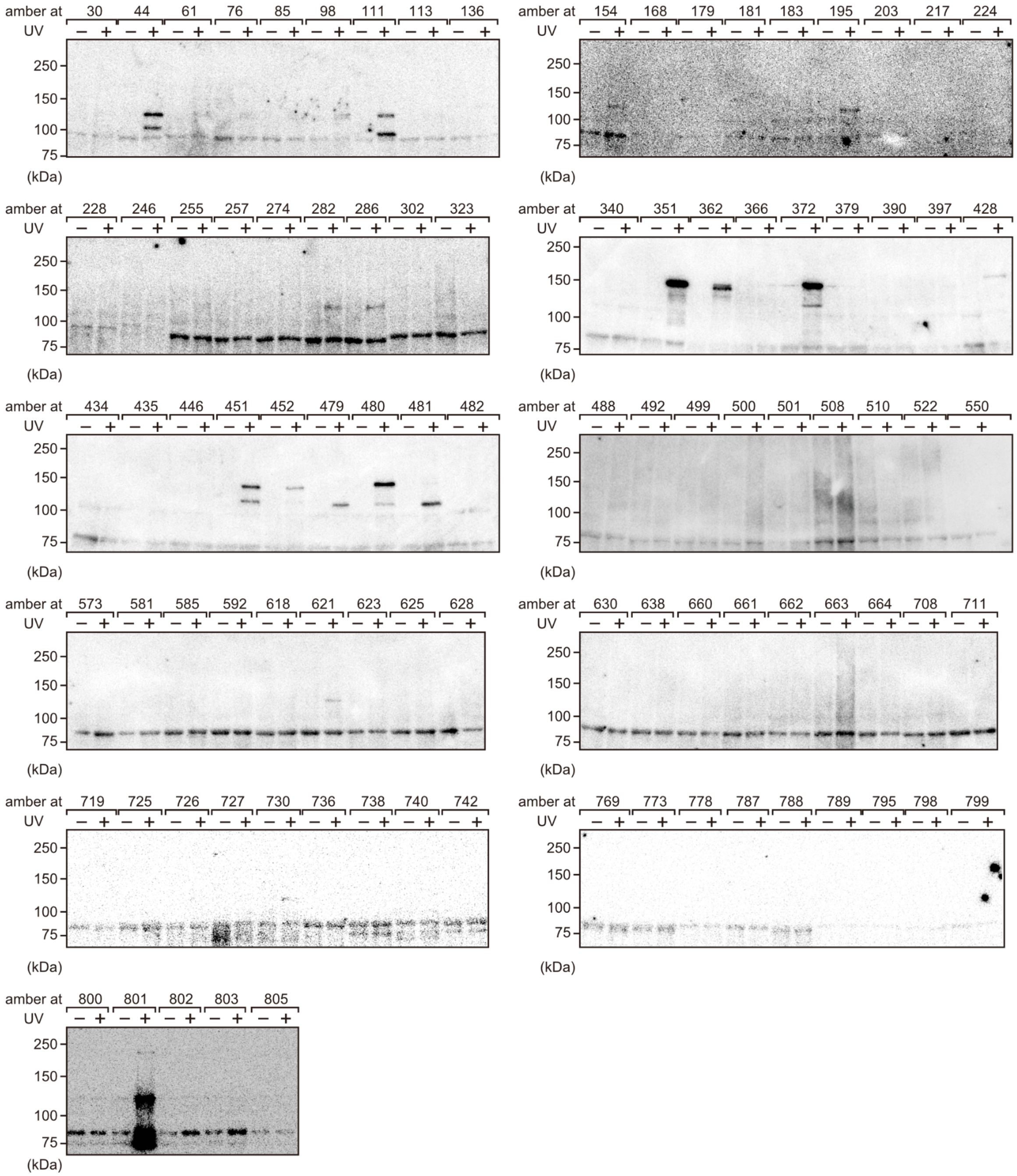


**Fig. S2.** The samples as in Fig. S1 were analyzed by immunoblotting as in Fig. S1 using anti-BamD antisera. BamA-BamD cross-linked products were identified at 44, 111, 154, 195, 282, 286, 351, 362, 372, 428, 451, 452, 480, 621, and 801 residues.

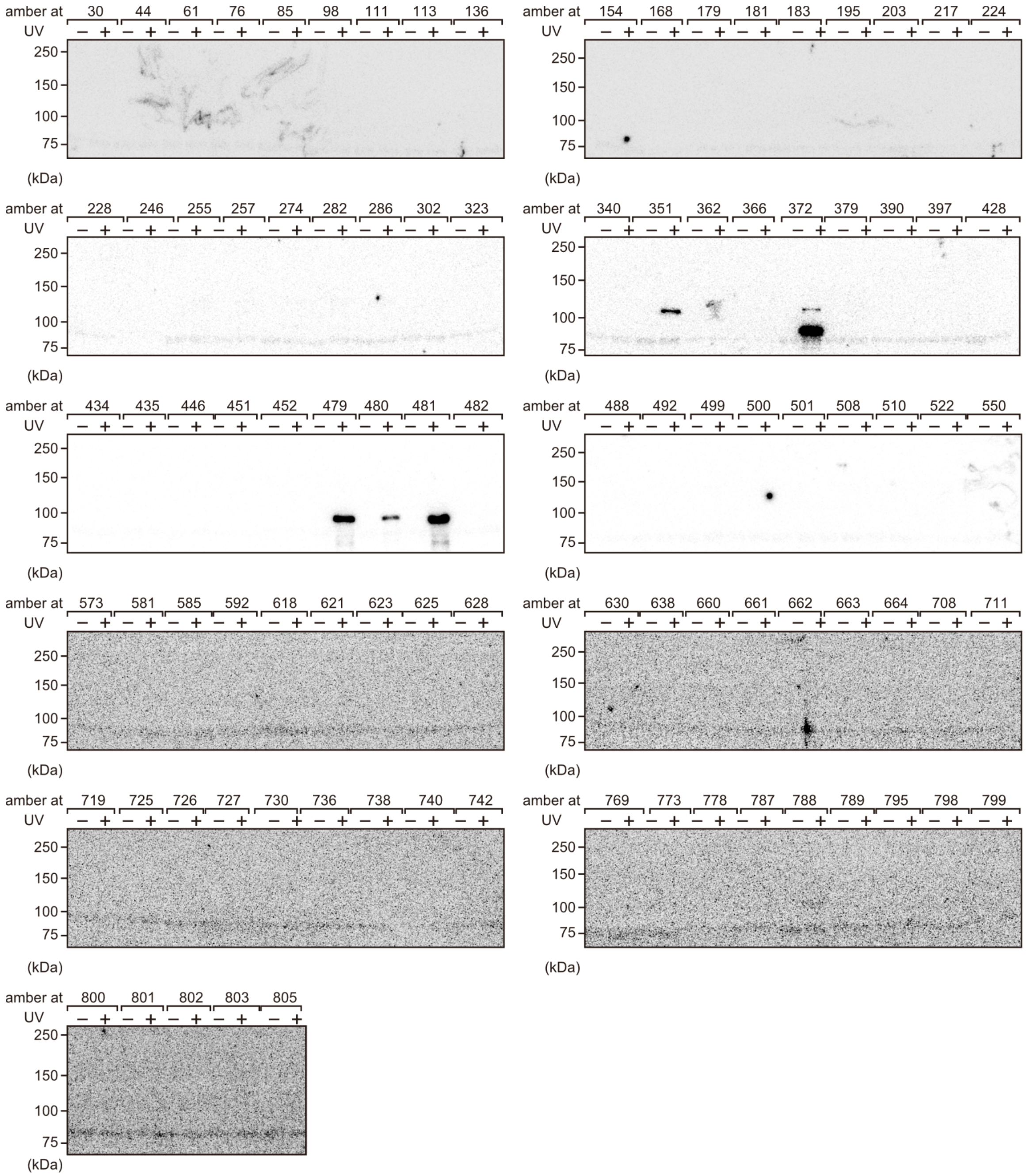

**Fig. S3.** The samples as in Fig. S1 were analyzed by immunoblotting as in Fig. S1 using anti-BamE antisera. BamA-BamE cross-linked products were identified at 351, 372, 479, 480, and 481 residues.

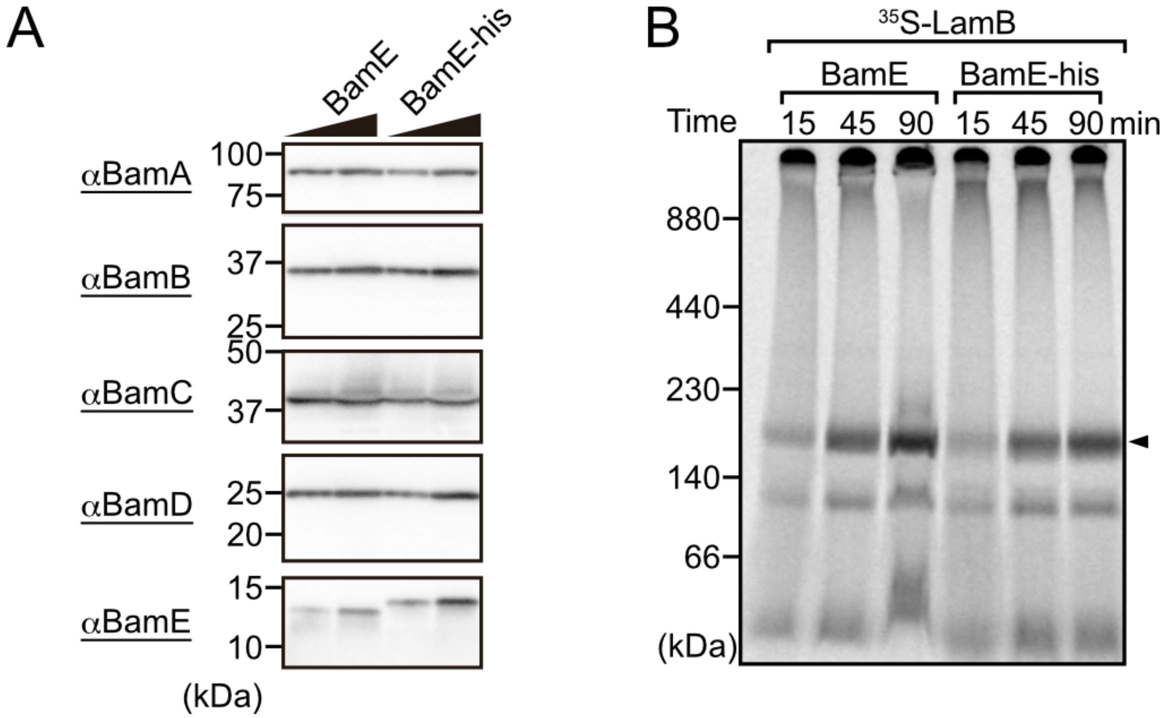


**Fig. S4.** The C-terminus attached octahistidine tag did not affect BamE function. (A) Comparing of the steady-state level of subunits of the BAM complex in BamE and BamE-his. Each subunit was analyzed by SDS-PAGE and immunoblotting using antisera recognizing the individual. (B) EMM isolated from indicated BamE expressing cell were incubated with radio-labeled LamB for indicated time at 30°C. EMM proteins were subjected to BN-PAGE and then analyzed by radio-imaging. Arrowhead indicates assembled trimeric-LamB.

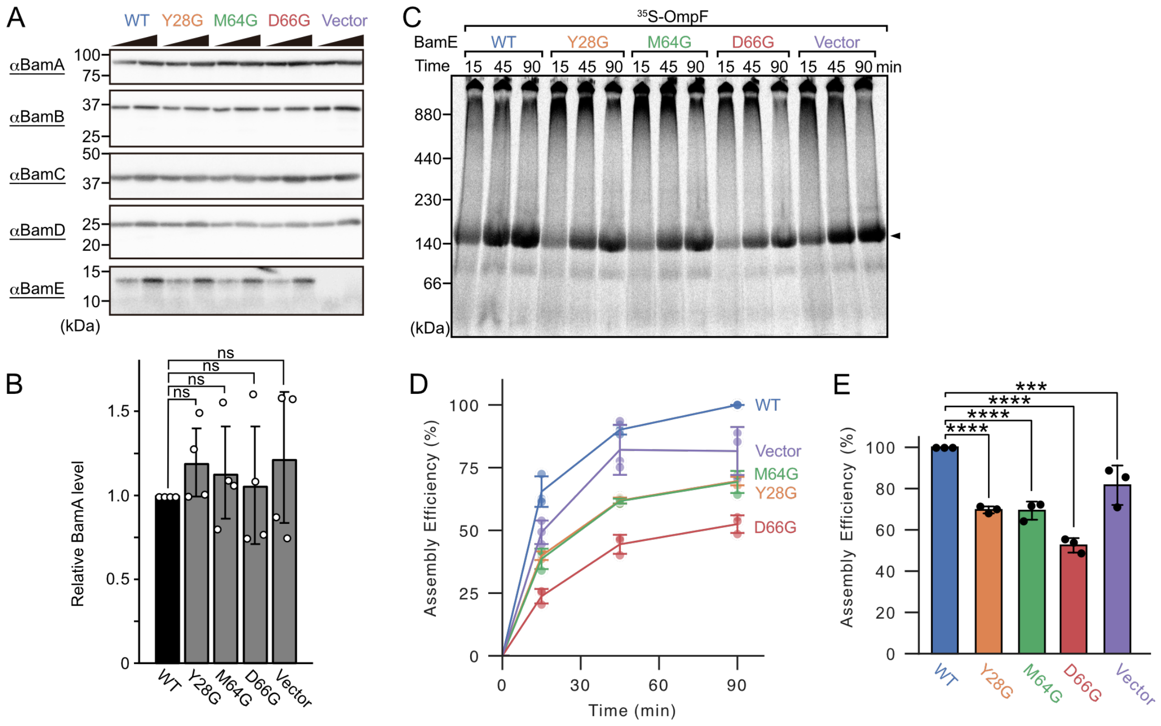


**Fig. S5.** BamE variants decrease OMPs assembly efficiency. (A) Steady-state level of all BAM complex components. Each subunit was analyzed by SDS-PAGE and immunoblotting using the antisera recognizing the individual. (B) Relative BamA level in BamE variants EMM. BamA level of WT set as control. Error bars represent standard deviation from four technical replicates. Statistical significance was indicated by signals as Figure 3 F (Dunnett's multiple comparison test). Exact values, WT vs Y28G: P=0.89, WT vs M64G: P=0.944, WT vs D66G: P=0.997, WT vs Vec: P=0.745. (C) EMM assembly assay using OmpF as a substrates as in supplementary figure. (D) Densitometry analysis of C. Amount of trimeric-OmpF at 90 minutes in WT set to 100%. Error bars represent standard deviation from three technical replicates. (E) Statistical analysis of assembly efficiency of trimeric-OmpF at 90 minutes. Error bars represent standard deviation from three technical replicates. Statistical significance was indicated by signals as Figure 3 F (Dunnett's multiple comparison test). Exact values, WT vs Y28G: P=7.61e-5, WT vs M64G: P=1.31e-4, WT vs D66G: P=2.74e-7, WT vs Vec: P=4.05e-3.

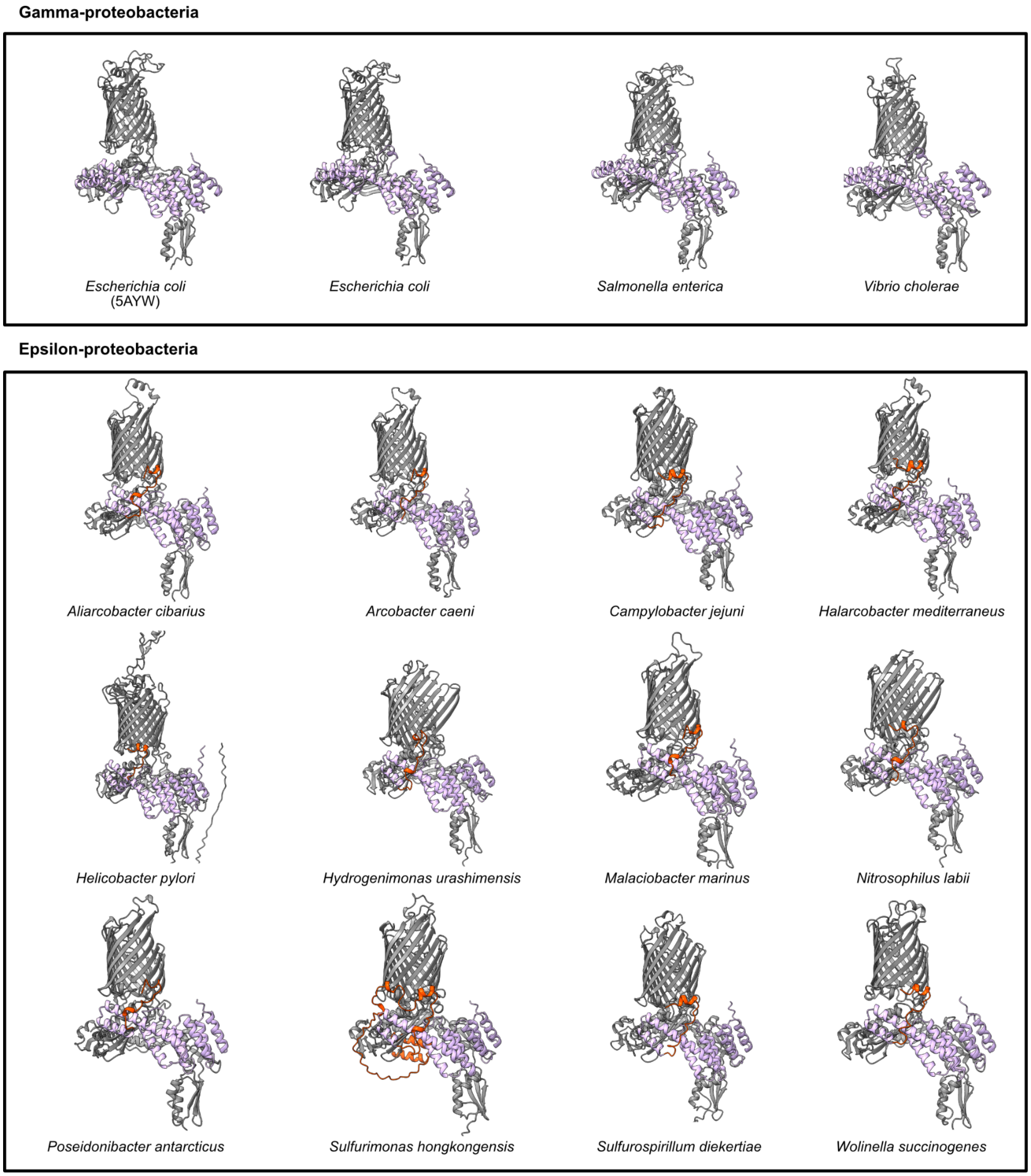


**Fig. S6.** C-terminal unstructured region of BamD in epsilon-proteobacteria is in close proximity to turn 1 and 2 of BamA. Structures of the BAM complex in gamma- (top) and epsilon-proteobacteria (bottom) are shown. The BAM complex of *Escherichia coli* determined by X-ray crystallography (5AYW) is shown on the left in the top panel. The other structures of the BAM complex in gamma-proteobacteria: *Escherichia coli*, *Salmonella enterica*, and *Vibrio cholerae*, and epsilon-proteobacteria: *Aliarcobacter cibarius*, *Arcobacter caeni*, *Campylobacter jejuni*, *Halarcobacter mediterraneus*, *Helicobacter pylori*, *Hydrogenimonas urashimensis*, *Malaciobacter marinus*, *Nitrosophilus labii*, *Poseidonibacter antarcticus*, *Sulfurimonas hongkongensis*, *Sulfurospirillum diekertiae*, and *Wolinella succinogenes*, are predicted using AlphaFold3. BamB, BamC, and BamE in gamma-proteobacteria are not shown. BamA and BamD are shown in gray and light purple, respectively. C-terminal unstructured region of BamD is highlighted in red.

**Supporting Tables**

**Table S1.** *Escherichia coli* strains used in this study.

| Strain | Genotype | Reference |
|---|---|---|
| BL21(DE3) * | $F^-$, *ompT*, hsdSB($r_B^-$ $m_B^-$), *gal*(λcI 857, *ind1*, *Sam7*, *nin5*, *lacUV5-T7gene1*), *dcm*(DE3) | Invitrogen |
| BamA shut-off | MC4100ara$^+$, Δ*bamA*::FRT *attB*::(*kan*, *araC*$^+$, $P_{BAD}$, *bamA*$^+$) | (1) |
| Δ*bamE* | BL21(DE3) *, *bamE*::*kan* | Lithgow laboratory |

**Table S2.** Plasmids used in this study.

| Plasmid name | Encoded gene and description | Primers for construct | Reference |
|---|---|---|---|
| pTnT | Expression vector, T7 polymerase promoter; $amp^R$ | -- | Promega |
| pTnT ($spe^R$) | Expression vector, T7 polymerase promoter; $spe^R$ | -- | This study |
| pSup-BpaRS-6TRN | The evolved *M. jannaschii* aminoacyl-tRNA synthetase / suppressor tRNA pair under control of the proK / a mutant glnS promoter; $Cm^R$ | -- | (2) |
| pEVOL-pBpF | The evolved *M. jannaschii* aminoacyl-tRNA synthetase / suppressor tRNA pair; $Cm^R$ | -- | (3) |
| pTnT-H6A2bamA (H30amb) | *bamA(H30amb)-his6* | -- | (4) |
| pTnT-H6A2bamA (L44amb) | *bamA(L44amb)-his6* | -- | (4) |
| pTnT-H6A2bamA (N61amb) | *bamA(N61amb)-his6* | -- | (4) |
| pTnT-H6A2bamA (R76amb) | *bamA(R76amb)-his6* | -- | (4) |
| pTnT-H6A2bamA (L85amb) | *bamA(L85amb)-his6* | -- | (4) |
| pTnT-H6A2bamA (T98amb) | *bamA(T98amb)-his6* | -- | (4) |
| pTnT-H6A2bamA (G111amb) | *bamA(G111amb)-his6* | -- | (4) |
| pTnT-H6A2bamA (K113amb) | *bamA(K113amb)-his6* | -- | (4) |
| pTnT-H6A2bamA (G136amb) | *bamA(G136amb)-his6* | -- | (4) |
| pTnT-H6A2bamA (V154amb) | *bamA(V154amb)-his6* | -- | (4) |
| pTnT-H6A2bamA (V168amb) | *bamA(V168amb)-his6* | -- | (4) |

| | | | |
|---|---|---|---|
| pTnT-H6A2bamA (Q179 amb) | *bamA(Q179amb)-his6* | -- | (5) |
| pTnT-H6A2bamA (N181amb) | *bamA(N181amb)-his6* | -- | (5) |
| pTnT-H6A2bamA (V183amb) | *bamA(V183amb)-his6* | -- | (5) |
| pTnT-H6A2bamA (S195amb) | *bamA(S195amb)-his6* | BamAS195amb-f BamAS195amb-r | This study |
| pTnT-H6A2bamA (V203amb) | *bamA(V203amb)-his6* | BamAV203amb-f BamAV203amb-r | This study |
| pTnT-H6A2bamA (Q217amb) | *bamA(217amb)-his6* | BamAQ217amb-f BamAQ217amb-r | This study |
| pTnT-H6A2bamA (E224amb) | *bamA(E224amb)-his6* | -- | (5) |
| pTnT-H6A2bamA (S228amb) | *bamA(S228amb)-his6* | BamAS228amb-f BamAS228amb-r | This study |
| pTnT-H6A2bamA (S246amb) | *bamA(S246amb)-his6* | -- | (5) |
| pTnT-H6A2bamA (Y255amb) | *bamA(Y255amb)-his6* | BamAY255amb-f BamAY255amb-r | This study |
| pTnT-H6A2bamA (T257amb) | *bamA(T257amb)-his6* | BamAT257amb-f BamAT257amb-r | This study |
| pTnT-H6A2bamA (S274amb) | *bamA(S274amb)-his6* | BamAS274amb-f BamAS274amb-r | This study |
| pTnT-H6A2bamA (A282amb) | *bamA(282amb)-his6* | BamAA282amb-f BamAA282amb-r | This study |
| pTnT-H6A2bamA (Q286amb) | *bamA(Q286amb)-his6* | BamAQ286amb-f BamAQ286amb-r | This study |
| pTnT-H6A2bamA (T302amb) | *bamA(T302amb)-his6* | BamAT302amb-f BamAT302amb-r | This study |
| pTnT-H6A2bamA (Q323amb) | *bamA(Q323amb)-his6* | BamAQ323amb-f BamAQ323amb-r | This study |
| pTnT-H6A2bamA (N340amb) | *bamA(N340amb)-his6* | BamAN340amb-f BamAN340amb-r | This study |
| pTnT-H6A2bamA (K351amb) | *bamA(K351amb)-his6* | BamAK351amb-f BamAK351amb-r | This study |

| | | | |
|---|---|---|---|
| pTnT-H6A2bamA (D362amb) | *bamA(D362amb)-his6* | BamAD362amb-f<br>BamAD362amb-r | This study |
| pTnT-H6A2bamA (R366amb) | *bamA(R366amb)-his6* | BamAR366amb-f<br>BamAR366amb-r | This study |
| pTnT-H6A2bamA (M372amb) | *bamA(M372amb)-his6* | BamAM372amb-f<br>BamAM372amb-r | This study |
| pTnT-H6A2bamA (S379amb) | *bamA(S379amb)-his6* | BamAS379amb-f<br>BamAS379amb-r | This study |
| pTnT-H6A2bamA (N390amb) | *bamA(N390amb)-his6* | BamAN390amb-f<br>BamAN390amb-r | This study |
| pTnT-H6A2bamA (T397amb) | *bamA(T397amb)-his6* | BamAT397amb-f<br>BamAT397amb-r | This study |
| pTnT-H6A2bamA (F428amb) | *bamA(F428amb)-his6* | BamAF428amb-f<br>BamAF428amb-r | This study |
| pTnT-H6A2bamA (T434amb) | *bamA(T434amb)-his6* | BamAT434amb-f<br>BamAT434amb-r | This study |
| pTnT-H6A2bamA (E435amb) | *bamA(E435amb)-his6* | BamAE435amb-f<br>BamAE435amb-r | This study |
| pTnT-H6A2bamA (Q446amb) | *bamA(Q446amb)-his6* | BamAQ445amb-f<br>BamAQ445amb-r | This study |
| pTnT-H6A2bamA (S451amb) | *bamA(S451amb)-his6* | BamAG451amb-f<br>BamAG451amb-r | This study |
| pTnT-H6A2bamA (M452amb) | *bamA(M452amb)-his6* | BamAT452amb-f<br>BamAT452amb-r | This study |
| pTnT-H6A2bamA (N479amb) | *bamA(N479amb)-his6* | BamAT479amb-f<br>BamAT479amb-r | This study |
| pTnT-H6A2bamA (V480amb) | *bamA(V480amb)-his6* | BamAV480amb-f<br>BamAV480amb-r | This study |
| pTnT-H6A2bamA (D481amb) | *bamA(D481amb)-his6* | BamAD481amb-f<br>BamAD481amb-r | This study |
| pTnT-H6A2bamA (G482amb) | *bamA(G482amb)-his6* | BamAG482amb-f<br>BamAG482amb-r | This study |
| pTnT-H6A2bamA (R488amb) | *bamA(R488amb)-his6* | BamAR488amb-f<br>BamAR488amb-r | This study |
| pTnT-H6A2bamA (N492amb) | *bamA(N492amb)-his6* | BamAN492amb-f<br>BamAN492amb-r | This study |

| | | | |
|---|---|---|---|
| pTnT-H6A2bamA (A499amb) | *bamA(A499amb)-his6* | BamAA499amb-f BamAA499amb-r | This study |
| pTnT-H6A2bamA (D500amb) | *bamA(D500amb)-his6* | BamAD500amb-f BamAD500amb-r | This study |
| pTnT-H6A2bamA (L501amb) | *bamA(L501amb)-his6* | BamAL501amb-f BamAL501amb-r | This study |
| pTnT-H6A2bamA (S508amb) | *bamA(S508amb)-his6* | BamAS508amb-f BamAS508amb-r | This study |
| pTnT-H6A2bamA (G510amb) | *bamA(G510amb)-his6* | BamAG510amb-f BamAG510amb-r | This study |
| pTnT-H6A2bamA (Y522amb) | *bamA Y522amb)-his6* | BamAY522amb-f BamAY522amb-r | This study |
| pTnT-H6A2bamA (Y550amb) | *bamA(Y550amb)-his6* | BamAY550amb-f BamAY550amb-r | This study |
| pTnT-H6A2bamA (N573amb) | *bamA(N573amb)-his6* | BamAN573amb-f BamAN573amb-r | This study |
| pTnT-H6A2bamA (L581amb) | *bamA(L581amb)-his6* | BamAL581amb-f BamAL581amb-r | This study |
| pTnT-H6A2bamA (Y585amb) | *bamA(Y585amb)-his6* | BamAY585amb-f BamAY585amb-r | This study |
| pTnT-H6A2bamA (R592amb) | *bamA(R592amb)-his6* | BamAR592amb-f BamAR592amb-r | This study |
| pTnT-H6A2bamA (Y618amb) | *bamA(Y618amb)-his6* | BamAY618amb-f BamAY618amb-r | This study |
| pTnT-H6A2bamA (I621amb) | *bamA(I621amb)-his6* | BamAI621amb-f BamAI621amb-r | This study |
| pTnT-H6A2bamA (D623amb) | *bamA(D623amb)-his6* | BamAD623amb-f BamAD623amb-r | This study |
| pTnT-H6A2bamA (H625amb) | *bamA(H625amb)-his6* | BamAH625amb-f BamAH625amb-r | This study |
| pTnT-H6A2bamA (V628amb) | *bamA(V628amb)-his6* | BamAV628amb-f BamAV628amb-r | This study |
| pTnT-H6A2bamA (L630amb) | *bamA(L630amb)-his6* | BamAL630amb-f BamAL630amb-r | This study |
| pTnT-H6A2bamA (G638amb) | *bamA(G638amb)-his6* | BamAG638amb-f BamAG638amb-r | This study |

| | | | |
|---|---|---|---|
| pTnT-H6A2bamA (V660amb) | *bamA(V660amb)-his6* | BamAV660amb-f BamAV660amb-r | This study |
| pTnT-H6A2bamA (R661amb) | *bamA(R661amb)-his6* | BamAR661amb-f BamAR661amb-r | This study |
| pTnT-H6A2bamA (G662amb) | *bamA(G662amb)-his6* | BamAG662amb-f BamAG662amb-r | This study |
| pTnT-H6A2bamA (F663amb) | *bamA(F663amb)-his6* | BamAF663amb-f BamAF663amb-r | This study |
| pTnT-H6A2bamA (Q664amb) | *bamA(Q664amb)-his6* | BamAQ664amb-f BamAQ664amb-r | This study |
| pTnT-H6A2bamA (G708amb) | *bamA(G708amb)-his6* | BamAG708amb-f BamAG708amb-r | This study |
| pTnT-H6A2bamA (M711amb) | *bamA(M711amb)-his6* | BamAM711amb-f BamAM711amb-r | This study |
| pTnT-H6A2bamA (I719amb) | *bamA(I719amb)-his6* | BamAI719amb-f BamAI719amb-r | This study |
| pTnT-H6A2bamA (I725amb) | *bamA(I725amb)-his6* | BamAI725amb-f BamAI725amb-r | This study |
| pTnT-H6A2bamA (S726amb) | *bamA(S726amb)-his6* | BamAS726amb-f BamAS726amb-r | This study |
| pTnT-H6A2bamA (D727amb) | *bamA(D727amb)-his6* | BamAD727amb-f BamAD727amb-r | This study |
| pTnT-H6A2bamA (A730amb) | *bamA(A730amb)-his6* | BamAA730amb-f BamAA730amb-r | This study |
| pTnT-H6A2bamA (S736amb) | *bamA(S736amb)-his6* | BamAS736amb-f BamAS736amb-r | This study |
| pTnT-H6A2bamA (F738amb) | *bamA(F738amb)-his6* | BamAF738amb-f BamAF738amb-r | This study |
| pTnT-H6A2bamA (D740amb) | *bamA(D740amb)-his6* | BamAD740amb-f BamAD740amb-r | This study |
| pTnT-H6A2bamA (G742amb) | *bamA(G742amb)-his6* | BamAG742amb-f BamAG742amb-r | This study |
| pTnT-H6A2bamA (S769amb) | *bamA(S769amb)-his6* | BamAS769amb-f BamAS769amb-r | This study |
| pTnT-H6A2bamA (A773amb) | *bamA(A773amb)-his6* | BamAA773amb-f BamAA773amb-r | This study |

| | | | |
|---|---|---|---|
| pTnT-H6A2bamA (S778amb) | *bamA(S778amb)-his6* | BamAS778amb-f BamAS778amb-r | This study |
| pTnT-H6A2bamA (Y787amb) | *bamA(Y787amb)-his6* | BamAY787amb-f BamAY787amb-r | This study |
| pTnT-H6A2bamA (A788amb) | *bamA(A788amb)-his6* | BamAA788amb-f BamAA788amb-r | This study |
| pTnT-H6A2bamA (Q789amb) | *bamA(Q789amb)-his6* | BamAQ789amb-f BamAQ789amb-r | This study |
| pTnT-H6A2bamA (D795amb) | *bamA(D795amb)-his6* | BamAD795amb-f BamAD795amb-r | This study |
| pTnT-H6A2bamA (K798amb) | *bamA(K798amb)-his6* | BamAK798amb-f BamAK798amb-r | This study |
| pTnT-H6A2bamA (A799amb) | *bamA(A799amb)-his6* | BamAA799amb-f BamAA799amb-r | This study |
| pTnT-H6A2bamA (E800amb) | *bamA(E800amb)-his6* | BamAE800amb-f BamAE800amb-r | This study |
| pTnT-H6A2bamA (Q801amb) | *bamA(Q801amb)-his6* | BamAQ801amb-f BamAQ801amb-r | This study |
| pTnT-H6A2bamA (F802amb) | *bamA(F802amb)-his6* | BamAF802amb-f BamAF802amb-r | This study |
| pTnT-H6A2bamA (Q803amb) | *bamA(Q803amb)-his6* | BamAQ803amb-f BamAQ803amb-r | This study |
| pTnT-H6A2bamA (N805amb) | *bamA(N805amb)-his6* | BamAN805amb-f BamAN805amb-r | This study |
| pTnT-bamE | *bamE* | BamE-f, BamE-r | This study |
| pTnT-bamE-his8 | *bamE-his8* | BamEhis8-f BamEhis8-r | This study |
| pTnT-bamE-his8 (P30amb) | *bamE(P30amb)-his8* | BamEP30amb-f BamEP30amb-r | This study |
| pTnT-bamE-his8 (N41amb) | *bamE(N41amb)-his8* | BamEN41amb-f BamEN41amb-r | This study |
| pTnT-bamE-his8 (V48amb) | *bamE(V48amb)-his8* | BamEV48amb-f BamEV48amb-r | This study |
| pTnT-bamE-his8 (Y57amb) | *bamE(Y57amb)-his8* | BamEY57amb-f BamEY57amb-r | This study |
| pTnT-bamE-his8 (L63amb) | *bamE(L63amb)-his8* | BamEL63amb-f BamEL63amb-r | This study |

| | | | |
|---|---|---|---|
| pTnT-bamE-his8 (M64amb) | *bamE(M64amb)-his8* | BamEM64amb-f BamEM64amb-r | This study |
| pTnT-bamE-his8 (F68amb) | *bamE(F68amb)-his8* | BamEF68amb-f BamEF68amb-r | This study |
| pTnT-bamE-his8 (G69amb) | *bamE(G69amb)-his8* | BamEQ69amb-f BamEQ69amb-r | This study |
| pTnT-bamE-his8 (T72amb) | *bamE(T72amb)-his8* | BamET72amb-f BamET72amb-r | This study |
| pTnT-bamE-his8 (F77amb) | *bamE(F77amb)-his8* | BamEF77amb-f BamEF77amb-r | This study |
| pTnT-bamE-his8 (R78amb) | *bamE(R78amb)-his8* | BamER78amb-f BamER78amb-r | This study |
| pTnT-bamE-his8 (H83amb) | *bamE(H83amb)-his8* | BamEH83amb-f BamEH83amb-r | This study |
| pTnT-bamE-his8 (V86amb) | *bamE(V86amb)-his8* | BamEV86amb-f BamEV86amb-r | This study |
| pTnT-bamE-his8 (T87amb) | *bamE(T87amb)-his8* | BamET87amb-f BamET87amb-r | This study |
| pTnT-bamE-his8 (T92amb) | *bamE(T92amb)-his8* | BamET92amb-f BamET92amb-r | This study |
| pTnT-bamE-his8 (S98amb) | *bamE(S98amb)-his8* | BamES98amb-f BamES98amb-r | This study |
| pTnT-bamE-his8 (N103amb) | *bamE(N103amb)-his8* | BamEN103amb-f BamEN103amb-r | This study |
| pTnT-bamE-his8 (N106amb) | *bamE(N106amb)-his8* | BamEN106amb-f BamEN106amb-r | This study |
| pTnT-bamE-his8 (K107amb) | *bamE(K107amb)-his8* | BamEK107amb-f BamEK107amb-r | This study |
| pTnT-bamE-his8 (Y28G) | *bamE(Y28G)-his8* | BamEY28G-f BamEY28G-r | This study |
| pTnT-bamE-his8 (M64G) | *bamE(M64G)-his8* | BamEM64G-f BamEM64G-r | This study |
| pTnT-bamE-his8 (D66G) | *bamE(D66G)-his8* | BamED66G-f BamED66G-r | This study |
| pTnT ($Spe^R$)-H6A2bamA (L44amb) | *bamA(L44amb)-his6* | BamAL44amb-f BamAL44amb-r | This study |

| | | | |
|---|---|---|---|
| pTnT (Spe$^R$)-H6A2bamA (K351amb) | *bamA(G111amb)-his6* | BamAK351amb-f BamAK351amb-r | This study |
| pTnT (Spe$^R$)-H6A2bamA (V480amb) | *bamA(M452amb)-his6* | BamAV480amb-f BamAV480amb-r | This study |
| pBAD-FLOmpC | *ompC* | -- | (6) |
| pBAD-FLOmpC (F280A Y286A) | *ompC FY* | -- | (6) |
| pTnT-LamB | *lamB* | -- | (6) |
| pTnT-OmpF | *ompF* | -- | (6) |

**Table S3.** Oligonucleotide primers used in this study.

| Primer name | Sequence5'->3' | Construction |
|---|---|---|
| BamAS195amb-f | ACCGACGAACTGATCtagCATTTCCAACTGCGT | Quick change |
| BamAS195amb-r | ACGCAGTTGGAAATGctaGATCAGTTCGTCGGT | Quick change |
| BamAV203amb-f | CAACTGCGTGACGAAtagCCGTGGTGGAACGTG | Quick change |
| BamAV203amb-r | CACGTTCCACCACGGctaTTCGTCACGCAGTTG | Quick change |
| BamAQ217amb-f | CGTAAATACCAGAAAtagAAACTGGCGGGCGAC | Quick change |
| BamAQ217amb-r | GTCGCCCGCCAGTTTctaTTTCTGGTATTTACG | Quick change |
| BamAS228amb-f | CTTGAAACCCTGCGCtagTACTATCTGGATCGC | Quick change |
| BamAS228amb-r | GCGATCCAGATAGTActaGCGCAGGGTTTCAAG | Quick change |
| BamAY255amb-f | GATAAAAAAGGTATTtagGTCACGGTGAACATC | Quick change |
| BamAY255amb-r | GATGTTCACCGTGACctaAATACCTTTTTTATC | Quick change |
| BamAT257amb-f | AAAGGTATTTACGTCtagGTGAACATCACCGAA | Quick change |
| BamAT257amb-r | TTCGGTGATGTTCACctaGACGTAAATACCTTT | Quick change |
| BamAS274amb-f | TCTGGCGTTGAAGTGtagGGCAACCTTGCCGGG | Quick change |
| BamAS274amb-r | CCCGGCAAGGTTGCCctaCACTTCAACGCCAGA | Quick change |
| BamAA282amb-f | CTTGCCGGGCACTCCtagGAAATTGAGCAGCTG | Quick change |
| BamAA282amb-r | CAGCTGCTCAATTTCctaGGAGTGCCCGGCAAG | Quick change |
| BamAQ286amb-f | TCCGCTGAAATTGAGtagCTGACTAAGATCGAG | Quick change |
| BamAQ286amb-r | CTCGATCTTAGTCAGctaCTCAATTTCAGCGGA | Quick change |
| BamAT302amb-f | AACGGCACCAAAGTGtagAAGATGGAAGATGAC | Quick change |
| BamAT302amb-r | GTCATCTTCCATCTTctaCACTTTGGTGCCGTT | Quick change |
| BamAQ323amb-f | GCCTATCCGCGCGTAtagTCGATGCCCGAAATT | Quick change |
| BamAQ323amb-r | AATTTCGGGCATCGActaTACGCGCGGATAGGC | Quick change |
| BamAN340amb-f | GTTAAATTACGTGTGtagGTTGATGCGGGTAAC | Quick change |
| BamAN340amb-r | GTTACCCGCATCAACctaCACACGTAATTTAAC | Quick change |
| BamAK351amb-f | CGTTTCTACGTGCGTtagATCCGTTTTGAAGGT | Quick change |
| BamAK351amb-r | ACCTTCAAAACGGATctaACGCACGTAGAAACG | Quick change |
| BamAD362amb-f | AACGATACCTCGAAAtagGCCGTCCTGCGTCGC | Quick change |
| BamAD362amb-r | GCGACGCAGGACGGCctaTTTCGAGGTATCGTT | Quick change |
| BamAR366amb-f | AAAGATGCCGTCCTGtagCGCGAAATGCGTCAG | Quick change |
| BamAR366amb-r | CTGACGCATTTCGCGctaCAGGACGGCATCTTT | Quick change |
| BamAM372amb-f | CGCGAAATGCGTCAGtagGAAGGTGCATGGCTG | Quick change |
| BamAM372amb-r | CAGCCATGCACCTTCctaCTGACGCATTTCGCG | Quick change |
| BamAS379amb-f | GGTGCATGGCTGGGGtagGATCTGGTCGATCAG | Quick change |
| BamAS379amb-r | CTGATCGACCAGATCctaCCCCAGCCATGCACC | Quick change |
| BamAN390amb-f | GGTAAGGAGCGTCTGtagCGTCTGGGCTTCTTT | Quick change |

| | | |
|---|---|---|
| BamAN390amb-r | AAAGAAGCCCAGACGctaCAGACGCTCCTTACC | Quick change |
| BamAT397amb-f | CTGGGCTTCTTTGAAtagGTCGATACCGATACC | Quick change |
| BamAT397amb-r | GGTATCGGTATCGACctaTTCAAAGAAGCCCAG | Quick change |
| BamAF428amb-f | ACCGGTAGCTTCAACtagGGTATTGGTTACGGT | Quick change |
| BamAF428amb-r | ACCGTAACCAATACCctaGTTGAAGCTACCGGT | Quick change |
| BamAT434amb-f | GGTATTGGTTACGGTtagGAAAGTGGCGTGAGC | Quick change |
| BamAT434amb-r | GCTCACGCCACTTTCctaACCGTAACCAATACC | Quick change |
| BamAE435amb-f | ATTGGTTACGGTACTtagAGTGGCGTGAGCTTC | Quick change |
| BamAE435amb-r | GAAGCTCACGCCACTctaAGTACCGTAACCAAT | Quick change |
| BamAQ446amb-f | CAGGCTGGTGTGCAGtagGATAACTGGTTAGGT | Quick change |
| BamAQ446amb-r | ACCTAACCAGTTATCctaCTGCACACCAGCCTG | Quick change |
| BamAG451amb-f | CAGGATAACTGGTTAtagACAGGTTATGCTGTT | Quick change |
| BamAG451amb-r | AACAGCATAACCTGTctaTAACCAGTTATCCTG | Quick change |
| BamAT452amb-f | GATAACTGGTTAGGTtagGGTTATGCTGTTGGT | Quick change |
| BamAT452amb-r | ACCAACAGCATAACCctaACCTAACCAGTTATC | Quick change |
| BamAT479amb-f | ACCAACCCGTACTTCtagGTAGATGGCGTAAGC | Quick change |
| BamAT479amb-r | GCTTACGCCATCTACctaGAAGTACGGGTTGGT | Quick change |
| BamAV480amb-f | AACCCGTACTTCACCtagGATGGCGTAAGCCTC | Quick change |
| BamAV480amb-r | GAGGCTTACGCCATCctaGGTGAAGTACGGGTT | Quick change |
| BamAD481amb-f | CCGTACTTCACCGTAtagGGCGTAAGCCTCGGT | Quick change |
| BamAD481amb-r | ACCGAGGCTTACGCCctaTACGGTGAAGTACGG | Quick change |
| BamAG482amb-f | TACTTCACCGTAGATtagGTAAGCCTCGGTGGT | Quick change |
| BamAG482amb-r | ACCACCGAGGCTTACctaATCTACGGTGAAGTA | Quick change |
| BamAR488amb-f | GTAAGCCTCGGTGGTtagCTCTTCTATAATGAC | Quick change |
| BamAR488amb-r | GTCATTATAGAAGAGctaACCACCGAGGCTTAC | Quick change |
| BamAN492amb-f | GGTCGTCTCTTCTATtagGACTTCCAGGCAGAT | Quick change |
| BamAN492amb-r | ATCTGCCTGGAAGTCctaATAGAAGAGACGACC | Quick change |
| BamAA499amb-f | TTCCAGGCAGATGACtagGACCTGTCCGACTAT | Quick change |
| BamAA499amb-r | ATAGTCGGACAGGTCctaGTCATCTGCCTGGAA | Quick change |
| BamAD500amb-f | CAGGCAGATGACGCCtagCTGTCCGACTATACC | Quick change |
| BamAD500amb-r | GGTATAGTCGGACAGctaGGCGTCATCTGCCTG | Quick change |
| BamAL501amb-f | GCAGATGACGCCGACtagTCCGACTATACCAAC | Quick change |
| BamAL501amb-r | GTTGGTATAGTCGGActaGTCGGCGTCATCTGC | Quick change |
| BamAS508amb-f | GACTATACCAACAAGtagTATGGTACAGACGTG | Quick change |
| BamAS508amb-r | CACGTCTGTACCATActaCTTGTTGGTATAGTC | Quick change |
| BamAG510amb-f | ACCAACAAGAGTTATtagACAGACGTGACGTTG | Quick change |
| BamAG510amb-r | CAACGTCACGTCTGTctaATAACTCTTGTTGGT | Quick change |

| | | |
|---|---|---|
| BamAY522amb-f | TTCCCGATTAACGAAtagAACTCGCTGCGTGCA | Quick change |
| BamAY522amb-r | TGCACGCAGCGAGTTctaTTCGTTAATCGGGAA | Quick change |
| BamAY550amb-f | ATGTGGCGTTATCTGtagTCTATGGGTGAACAT | Quick change |
| BamAY550amb-r | ATGTTCACCCATAGActaCAGATAACGCCACAT | Quick change |
| BamAN573amb-f | GACGACTTCACGTTCtagTATGGTTGGACCTAT | Quick change |
| BamAN573amb-r | ATAGGTCCAACCATActaGAACGTGAAGTCGTC | Quick change |
| BamAL581amb-f | TGGACCTATAACAAGtagGACCGTGGTTACTTC | Quick change |
| BamAL581amb-r | GAAGTAACCACGGTCctaCTTGTTATAGGTCCA | Quick change |
| BamAY585amb-f | AAGCTTGACCGTGGTtagTTCCCGACAGATGGT | Quick change |
| BamAY585amb-r | ACCATCTGTCGGGAActaACCACGGTCAAGCTT | Quick change |
| BamAR592amb-f | CCGACAGATGGTTCAtagGTCAACCTGACCGGT | Quick change |
| BamAR592amb-r | ACCGGTCAGGTTGACctaTGAACCATCTGTCGG | Quick change |
| BamAY618amb-f | TTAGACACGGCGACTtagGTGCCGATCGATGAC | Quick change |
| BamAY618amb-r | GTCATCGATCGGCACctaAGTCGCCGTGTCTAA | Quick change |
| BamAI621amb-f | GCGACTTATGTGCCGtagGATGACGATCACAAA | Quick change |
| BamAI621amb-r | TTTGTGATCGTCATCctaCGGCACATAAGTCGC | Quick change |
| BamAD623amb-f | TATGTGCCGATCGATtagGATCACAAATGGGTT | Quick change |
| BamAD623amb-r | AACCCATTTGTGATCctaATCGATCGGCACATA | Quick change |
| BamAH625amb-f | CCGATCGATGACGATtagAAATGGGTTGTTCTG | Quick change |
| BamAH625amb-r | CAGAACAACCCATTTctaATCGTCATCGATCGG | Quick change |
| BamAV628amb-f | GACGATCACAAATGGtagGTTCTGGGGCGTACC | Quick change |
| BamAV628amb-r | GGTACGCCCCAGAACctaCCATTTGTGATCGTC | Quick change |
| BamAL630amb-f | CACAAATGGGTTGTTtagGGGCGTACCCGCTGG | Quick change |
| BamAL630amb-r | CCAGCGGGTACGCCCctaAACAACCCATTTGTG | Quick change |
| BamAG638amb-f | ACCCGCTGGGGTTATtagGATGGTTTAGGCGGC | Quick change |
| BamAG638amb-r | GCCGCCTAAACCATCctaATAACCCCAGCGGGT | Quick change |
| BamAV660amb-f | GGTGGTTCCAGCACCtagCGTGGCTTCCAGTCC | Quick change |
| BamAV660amb-r | GGACTGGAAGCCACGctaGGTGCTGGAACCACC | Quick change |
| BamAR661amb-f | GGTTCCAGCACCGTGtagGGCTTCCAGTCCAAT | Quick change |
| BamAR661amb-r | ATTGGACTGGAAGCCctaCACGGTGCTGGAACC | Quick change |
| BamAG662amb-f | TCCAGCACCGTGCGTtagTTCCAGTCCAATACC | Quick change |
| BamAG662amb-r | GGTATTGGACTGGAActaACGCACGGTGCTGGA | Quick change |
| BamAF663amb-f | AGCACCGTGCGTGGCtagCAGTCCAATACCATT | Quick change |
| BamAF663amb-r | AATGGTATTGGACTGctaGCCACGCACGGTGCT | Quick change |
| BamAQ664amb-f | ACCGTGCGTGGCTTCtagTCCAATACCATTGGT | Quick change |
| BamAQ664amb-r | ACCAATGGTATTGGActaGAAGCCACGCACGGT | Quick change |
| BamAG708amb-f | GATGATGCTGTAGGCtagAACGCCATGGCGGTT | Quick change |

| | | |
|---|---|---|
| BamAG708amb-r | AACCGCCATGGCGTTctaGCCTACAGCATCATC | Quick change |
| BamAM711amb-f | GTAGGCGGTAACGCCtagGCGGTTGCCAGCCTC | Quick change |
| BamAM711amb-r | GAGGCTGGCAACCGCctaGGCGTTACCGCCTAC | Quick change |
| BamAI719amb-f | GCCAGCCTCGAGTTCtagACCCCGACGCCGTTT | Quick change |
| BamAI719amb-r | AAACGGCGTCGGGGTctaGAACTCGAGGCTGGC | Quick change |
| BamAI725amb-f | ACCCCGACGCCGTTTtagAGCGATAAGTATGCT | Quick change |
| BamAI725amb-r | AGCATACTTATCGCTctaAAACGGCGTCGGGGT | Quick change |
| BamAS726amb-f | CCGACGCCGTTTATTtagGATAAGTATGCTAAC | Quick change |
| BamAS726amb-r | GTTAGCATACTTATCctaAATAAACGGCGTCGG | Quick change |
| BamAD727amb-f | ACGCCGTTTATTAGCtagAAGTATGCTAACTCG | Quick change |
| BamAD727amb-r | CGAGTTAGCATACTTctaGCTAATAAACGGCGT | Quick change |
| BamAA730amb-f | ATTAGCGATAAGTATtagAACTCGGTTCGTACT | Quick change |
| BamAA730amb-r | AGTACGAACCGAGTTctaATACTTATCGCTAAT | Quick change |
| BamAS736amb-f | AACTCGGTTCGTACTtagTTCTTCTGGGATATG | Quick change |
| BamAS736amb-r | CATATCCCAGAAGAActaAGTACGAACCGAGTT | Quick change |
| BamAF738amb-f | GTTCGTACTTCCTTCtagTGGGATATGGGTACC | Quick change |
| BamAF738amb-r | GGTACCCATATCCCActaGAAGGAAGTACGAAC | Quick change |
| BamAD740amb-f | ACTTCCTTCTTCTGGtagATGGGTACCGTTTGG | Quick change |
| BamAD740amb-r | CCAAACGGTACCCATctaCCAGAAGAAGGAAGT | Quick change |
| BamAG742amb-f | TTCTTCTGGGATATGtagACCGTTTGGGATACA | Quick change |
| BamAG742amb-r | TGTATCCCAAACGGTctaCATATCCCAGAAGAA | Quick change |
| BamAS769amb-f | AGCAATATCCGTATGtagGCGGGTATCGCATTA | Quick change |
| BamAS769amb-r | TAATGCGATACCCGCctaCATACGGATATTGCT | Quick change |
| BamAA773amb-f | ATGTCTGCGGGTATCtagTTACAATGGATGTCC | Quick change |
| BamAA773amb-r | GGACATCCATTGTAActaGATACCCGCAGACAT | Quick change |
| BamAS778amb-f | GCATTACAATGGATGtagCCATTGGGGCCGTTG | Quick change |
| BamAS778amb-r | CAACGGCCCCAATGGctaCATCCATTGTAATGC | Quick change |
| BamAY787amb-f | CCGTTGGTGTTCTCCtagGCCCAGCCGTTCAAA | Quick change |
| BamAY787amb-r | TTTGAACGGCTGGGCctaGGAGAACACCAACGG | Quick change |
| BamAA788amb-f | TTGGTGTTCTCCTACtagCAGCCGTTCAAAAAG | Quick change |
| BamAA788amb-r | CTTTTTGAACGGCTGctaGTAGGAGAACACCAA | Quick change |
| BamAQ789amb-f | GTGTTCTCCTACGCCtagCCGTTCAAAAAGTAC | Quick change |
| BamAQ789amb-r | GTACTTTTTGAACGGctaGGCGTAGGAGAACAC | Quick change |
| BamAD795amb-f | CCGTTCAAAAAGTACtagTGGAGACAAGGCAGA | Quick change |
| BamAD795amb-r | TCTGCCTTGTCTCCActaGTACTTTTTGAACGG | Quick change |
| BamAK798amb-f | AAGTACGATGGAGACtagGCAGAACAGTTCCAG | Quick change |
| BamAK798amb-r | CTGGAACTGTTCTGCctaGTCTCCATCGTACTT | Quick change |

| | | |
|---|---|---|
| BamAA799amb-f | TACGATGGAGACAAGtagGAACAGTTCCAGTTT | Quick change |
| BamAA799amb-r | AAACTGGAACTGTTCctaCTTGTCTCCATCGTA | Quick change |
| BamAE800amb-f | GATGGAGACAAGGCAtagCAGTTCCAGTTTAAC | Quick change |
| BamAE800amb-r | GTTAAACTGGAACTGctaTGCCTTGTCTCCATC | Quick change |
| BamAQ801amb-f | GGAGACAAGGCAGAAtagTTCCAGTTTAACATC | Quick change |
| BamAQ801amb-r | GATGTTAAACTGGAActaTTCTGCCTTGTCTCC | Quick change |
| BamAF802amb-f | GACAAGGCAGAACAGtagCAGTTTAACATCGGT | Quick change |
| BamAF802amb-r | ACCGATGTTAAACTGctaCTGTTCTGCCTTGTC | Quick change |
| BamAQ803amb-f | AAGGCAGAACAGTTCtagTTTAACATCGGTAAA | Quick change |
| BamAQ803amb-r | TTTACCGATGTTAAActaGAACTGTTCTGCCTT | Quick change |
| BamAN805amb-f | GAACAGTTCCAGTTTtagATCGGTAAAACCTGG | Quick change |
| BamAN805amb-r | CCAGGTTTTACCGATctaAAACTGGAACTGTTC | Quick change |
| BamE-f | aacgcataataacgtctagaATGCGCTGTAAAACGCTGACTGC TGCAGCAGCAGTACTAT | SLiCE |
| BamE-r | ttgcggccgcccgggtcgacTTGAGCACCTTTTTAACGTCTTTGAGA GCAACTTTATTA | SLiCE |
| BamEhis8-f | CCTGCGCTGAGTGGTAACCATCATCACCATCACCATC ACCATTAATAATAAAGTTGCTCT | Quick change |
| BamEhis8-r | AGAGCAACTTTATTATTAATGGTGATGGTGATGGTGAT GATGGTTACCACTCAGCGCAGG | Quick change |
| BamEP30amb-f | GCGAGTGGTTTACCGTtagGACATCAACCAGGGGA | Quick change |
| BamEP30amb-r | TCCCCTGGTTGATGTCctaACGGTAAACCACTCGC | Quick change |
| BamEN41amb-f | GAACTATCTGACCGCTtagGACGTATCCAAAATAC | Quick change |
| BamEN41amb-r | GTATTTTGGATACGTCctaAGCGGTCAGATAGTTC | Quick change |
| BamEV48amb-f | CGTATCCAAAATACGTtagGGCATGACGCAACAAC | Quick change |
| BamEV48amb-r | GTTGTTGCGTCATGCCctaACGTATTTTGGATACG | Quick change |
| BamEY57amb-f | GCAACAACAAGTTGCGtagGCATTGGGTACACCGC | Quick change |
| BamEY57amb-r | GCGGTGTACCCAATGCctaCGCAACTTGTTGTTGC | Quick change |
| BamEL63amb-f | CGCATTGGGTACACCGtagATGTCCGATCCATTTG | Quick change |
| BamEL63amb-r | CAAATGGATCGGACATctaCGGTGTACCCAATGCG | Quick change |
| BamEM64amb-f | ATTGGGTACACCGCTGtagTCCGATCCATTTGGTA | Quick change |
| BamEM64amb-r | TACCAAATGGATCGGActaCAGCGGTGTACCCAAT | Quick change |
| BamEF68amb-f | GCTGATGTCCGATCCAtagGGTACGAATACCTGGT | Quick change |
| BamEF68amb-r | ACCAGGTATTCGTACCctaTGGATCGGACATCAGC | Quick change |
| BamEQ69amb-f | GATGTCCGATCCATTTtagACGAATACCTGGTTCT | Quick change |
| BamEQ69amb-r | AGAACCAGGTATTCGTctaAAATGGATCGGACATC | Quick change |
| BamEN71amb-f | CGATCCATTTGGTACGtagACCTGGTTCTATGTCT | Quick change |
| BamRN71amb-r | AGACATAGAACCAGGTctaCGTACCAAATGGATCG | Quick change |

| | | |
|---|---|---|
| BamET72amb-f | TCCATTTGGTACGAATtagTGGTTCTATGTCTTCC | Quick change |
| BamET72amb-r | GGAAGACATAGAACCActaATTCGTACCAAATGGA | Quick change |
| BamEF77amb-f | TACCTGGTTCTATGTCtagCGCCAGCAACCAGGTC | Quick change |
| BamEF77amb-r | GACCTGGTTGCTGGCGctaGACATAGAACCAGGTA | Quick change |
| BamER78amb-f | CTGGTTCTATGTCTTCtagCAGCAACCAGGTCATG | Quick change |
| BamER78amb-r | CATGACCTGGTTGCTGctaGAAGACATAGAACCAG | Quick change |
| BamEH83amb-f | CCGCCAGCAACCAGGTtagGAAGGTGTAACTCAGC | Quick change |
| BamEH83amb-r | GCTGAGTTACACCTTCctaACCTGGTTGCTGGCGG | Quick change |
| BamEV86amb-f | ACCAGGTCATGAAGGTtagACTCAGCAAACGCTGA | Quick change |
| BamEV86amb-r | TCAGCGTTTGCTGAGTctaACCTTCATGACCTGGT | Quick change |
| BamET87amb-f | AGGTCATGAAGGTGTAtagCAGCAAACGCTGACGC | Quick change |
| BamET87amb-r | GCGTCAGCGTTTGCTGctaTACACCTTCATGACCT | Quick change |
| BamET92amb-f | AACTCAGCAAACGCTGtagCTGACCTTTAACAGTA | Quick change |
| BamET92amb-r | TACTGTTAAAGGTCAGctaCAGCGTTTGCTGAGTT | Quick change |
| BamES98amb-f | GCTGACCTTTAACAGTtagGGTGTGTTGACCAATA | Quick change |
| BamES98amb-r | TATTGGTCAACACACCctaACTGTTAAAGGTCAGC | Quick change |
| BamEN103amb-f | TAGCGGTGTGTTGACCtagATTGATAACAAACCTG | Quick change |
| BamEN103amb-r | CAGGTTTGTTATCAATctaGGTCAACACACCGCTA | Quick change |
| BamEN106amb-f | GTTGACCAATATTGATtagAAACCTGCGCTGAGTG | Quick change |
| BamEN106amb-r | CACTCAGCGCAGGTTTctaATCAATATTGGTCAAC | Quick change |
| BamEK107amb-f | GACCAATATTGATAACtagCCTGCGCTGAGTGGTA | Quick change |
| BamEK107amb-r | TACCACTCAGCGCAGGctaGTTATCAATATTGGTC | Quick change |
| BamEY28G-f | CTGGAGCGAGTGGTTggcCGTCCTGACATCAAC | Quick change |
| BamEY28G-r | GTTGATGTCAGGACGgccAACCACTCGCTCCAG | Quick change |
| BamEM64G-f | TTGGGTACACCGCTGggcTCCGATCCATTTGGT | Quick change |
| BamEM64G-r | ACCAAATGGATCGGAgccCAGCGGTGTACCCAA | Quick change |
| BamED66G-f | ACACCGCTGATGTCCggcCCATTTGGTACGAAT | Quick change |
| BamED66G-r | ATTCGTACCAAATGGgccGGACATCAGCGGTGT | Quick change |
| pET-Spe$^R$1 | agtaaacttggtctgacagttatttgccgactaccttggtgatctcgcc | SLiCE |
| pET-Spe$^R$2 | atattgaaaaaggaagagtatgagggaagcggtgatcgccgaagtatcg | SLiCE |
| pET-Spe$^R$3 | actcttcctttttcaatattattgaagcatttatc | SLiCE |
| pET-Spe$^R$4 | ctgtcagaccaagtttactcatatatactttagat | SLiCE |

**Table S4.** Sequence of synthesized spectinomycin resistant gene

5’ -
TTATTTGCCGACTACCTTGGTGATCTCGCCTTTCACGTAGTGGACAAATTCTTCCAACTGATC
TGCGCGCGAGGCCAGCGATCTTCTTCTGTCCAAGATAAGCCTGTCTAGCTCAGTATGACGG
GCTGATACTGGGCCGGCAGGCGCTCCATTGCCCAGTCGGCAGCGACATCCTTCGGCGCGA
TTTTGCCGGTTACTGCGCTGTACCAAATGCGGGACAACGTAAGCACTACATTTCGCTCATCG
CCAGCCCAGTCGGGCGGCGAGTTCCATAGCGTTAAGGTTTCATTTAGCGCCTCAAATAGAT
CCTGTTCAGGAACCGGATCAAAGAGTTCCTCCGCCGCTGGACCTACCAAGGCAACGCTATG
TTCTCTTGCTTTTGTCAGCAAGATAGCCAGATCAATGTCGATCGTGGCTGGCTCGAAGATAC
CTGCAAGAATGTCATTGCGCTGCCATTCTCCAAATTGCAGTTCGCGCTTAGCTGGATAACGC
CACGGAATGATGTCGTCGTGCACAACAATGGTGACTTCTACAGCGCGGAGAATCTCGCTCT
CTCCAGGGGAAGCCGAAGTTTCCAAAAGGTCGTTGATCAAAGCTCGCCGCGTTGTTTCATCA
AGCCTTACGGTCACCGTAACCAGCAAATCAATATCACTGTGTGGCTTCAGGCCGCCATCCAC
TGCGGAGCCGTACAAATGTACGGCCAGCAACGTCGGTTCGAGATGGCGCTCGATGACGCC
AACTACCTCTGATAGTTGAGTCGATACTTCGGCGATCACCGCTTCCCTCAT - 3’

**SI References**


1. U. Lehr, *et al.*, C-terminal amino acid residues of the trimeric autotransporter adhesin YadA of Yersinia enterocolitica are decisive for its recognition and assembly by BamA: YadA is recognized and assembled by the BAM complex. *Mol. Microbiol.* **78**, 932–946 (2010).

2. Y. Ryu, P. G. Schultz, Efficient incorporation of unnatural amino acids into proteins in Escherichia coli. *Nat. Methods* **3**, 263–265 (2006).

3. J. W. Chin, P. G. Schultz, In Vivo Photocrosslinking with Unnatural Amino Acid Mutagenesis. *ChemBioChem* **3**, 1135–1137 (2002).

4. S. D. Gunasinghe, *et al.*, The WD40 Protein BamB Mediates Coupling of BAM Complexes into Assembly Precincts in the Bacterial Outer Membrane. *Cell Rep.* **23**, 2782–2794 (2018).

5. Y. Daimon, *et al.*, The TPR domain of BepA is required for productive interaction with substrate proteins and the β-barrel assembly machinery complex. *Mol. Microbiol.* **106**, 760–776 (2017).

6. E. M. Germany, *et al.*, Dual recognition of multiple signals in bacterial outer membrane proteins enhances assembly and maintains membrane integrity. *eLife* **12**, RP90274 (2024).
